# Flp-recombinase mouse line for genetic manipulation of ipRGCs

**DOI:** 10.1101/2024.05.06.592761

**Authors:** E Contreras, C Liang, HL Mahoney, JL Javier, ML Luce, K Labastida Medina, T Bozza, TM Schmidt

## Abstract

Light has myriad impacts on behavior, health, and physiology. These signals originate in the retina and are relayed to the brain by more than 40 types of retinal ganglion cells (RGCs). Despite a growing appreciation for the diversity of RGCs, how these diverse channels of light information are ultimately integrated by the ∼50 retinorecipient brain targets to drive these light-evoked effects is a major open question. This gap in understanding primarily stems from a lack of genetic tools that specifically label, manipulate, or ablate specific RGC types. Here, we report the generation and characterization of a new mouse line (Opn4^FlpO^), in which FlpO is expressed from the *Opn4* locus, to manipulate the melanopsin-expressing, intrinsically photosensitive retinal ganglion cells. We find that the Opn4^FlpO^ line, when crossed to multiple reporters, drives expression that is confined to ipRGCs and primarily labels the M1-M3 subtypes. Labeled cells in this mouse line show the expected intrinsic, melanopsin-based light response and morphological features consistent with the M1-M3 subtypes. In alignment with the morphological and physiological findings, we see strong innervation of non-image forming brain targets by ipRGC axons, and weaker innervation of image forming targets in Opn4^FlpO^ mice labeled using AAV-based and FlpO-reporter lines. Consistent with the FlpO insertion disrupting the endogenous Opn4 transcript, we find that Opn4^FlpO/FlpO^ mice show deficits in the pupillary light reflex, demonstrating their utility for behavioral research in future experiments. Overall, the Opn4^FlpO^ mouse line drives Flp-recombinase expression that is confined to ipRGCs and most effectively drives recombination in M1-M3 ipRGCs. This mouse line will be of broad use to those interested in manipulating ipRGCs through a Flp-based recombinase for intersectional studies or in combination with other, non-Opn4 Cre driver lines.

## Introduction

Light has broad impacts on behavior beyond conscious visual perception, including entraining our daily activity, sleep, and physiological rhythms to the light/dark cycle, altering pupil diameter in response to changes in environmental light, and even affecting our mood and ability to learn (Hattar et al., 2003; Mrosovsky and Hattar, 2003; Lucas et al., 2003; Panda et al., 2003; Gooley et al., 2012; LeGates et al., 2012; Huang, Lu et al. 2012; Fernandez et al., 2018; Schwartz, 2015; Wirz-Justice et al., 2013; Mahoney and Schmidt, 2024). Each of these behaviors initiates with the capture of light by photoreceptors in the retina. In the mouse, there are more than 50 types of retinal ganglion cells (RGCs) (Morin and Studholme, 2014; Martersteck et al., 2017). Each type relays information about different features of the visual scene to a specific subset of 50 downstream retinorecipient targets in the brain to impact specific behaviors (Morin and Studholme, 2014; Martersteck et al., 2017). Despite a growing appreciation of the stunning diversity of cell types and circuits driving these broad behavioral impacts of light, how the signals and circuits of individual RGC types impact distinct behaviors is poorly understood. This lack of understanding has arisen in part due to the paucity of genetic tools to specifically label, manipulate, or ablate RGC subtypes. Such manipulations often rely on genetically modified mice in which Cre recombinase is expressed in specific neuronal populations. While such “Cre-driver” lines exist that differentially target RGCs, expression is often not confined solely to a single RGC type, and thus may tag multiple populations of RGCs or even other retinal neuron types.

The intrinsically photosensitive (ip)RGCs represent one of the most genetically tractable RGCs due to the specific expression of the gene *Opn4*, which encodes the light sensitive protein melanopsin (Berson et al., 2002; Hattar et al., 2002; Provencio et al., 2002; Gong et., 2003; Schmidt et al., 2008; Do et al., 2009; Güler et a., 2008; Ecker et al., 2010; Berson et al., 2010; Chen et al., 2011; Reifler et al., 2023). Because of its specific expression pattern, *Opn4* has been used to target expression of Cre selectively in ipRGCs to manipulate gene expression through crossing with Cre-dependent mouse lines or transduction with Cre-dependent AAV vectors (Ecker et al., 2010; Chen et al., 2011; Delwig et al., 2016; Reifler et al., 2023; Aranda and Schmidt, 2021). These genetic tools have helped reveal critical roles of ipRGCs in circadian photoentrainment, sleep, contrast sensitivity for visual perception, the pupillary light reflex, learning, and mood (Hattar et al., 2002, 2006; Gooley et al., 2003; Baver et al., 2008; Panda et al., 2002; Ruby et al., 2002; Hatori et al., 2008; Göz et al., 2008; Güler et al., 2008; Chew et al., 2017; Lucas et al., 2003; Altimus et al., 2008; Lupi et al., 2008; Rupp et al., 2019; Chen et al., 2011; LeGates et al., 2012; Fernandez et al., 2018; Schmidt et al., 2014). However, there are in fact six subtypes of ipRGC (M1-M6), each of which possesses a distinct subset of morphological and physiological properties, central projections, and behavioral roles (Sondereker et al., 2021; Aranda and Schmidt, 2021; Contreras et al., 2021; Mahoney and Schmidt, 2024). Thus, even for this highly genetically tractable RGC population, existing tools do not allow us to specifically label, manipulate, or ablate single ipRGC subtypes. Thus, how each of these very different subtypes might contribute to the wide variety of ipRGC-driven behaviors is still an open question.

It is becoming increasingly clear that targeting Cre expression under the control of single genes will not be specific sufficient to tackle the fundamental question of how individual retinal cell types contribute to light-evoked behaviors. One potential solution is the use of intersectional genetic approaches that increase specificity by selectively marking cells that express both Cre-recombinase and FlpO-recombinase (Awatramani et al., 2001; Farago et al., 2006; Miyoshi and Fishell, 2006; Sousa et al, 2009; Yamamoto et al, 2009; Imayoshi et al., 2012; Daigle et al., 2018; Weinholtz and Castle, 2021). Recent years have seen an explosion of intersectional Cre and FlpO-dependent reporter mouse lines and AAV reporter vectors that exhibit genetic activity in the presence of both recombinases, allowing highly specific cell-type and circuit-based manipulations (Awatramani et al., 2001; Farago et al., 2006; Miyoshi and Fishell, 2006; Sousa et al, 2009; Yamamoto et al, 2009; Imayoshi et al., 2012; Daigle et al., 2018; Weinholtz and Castle, 2021). There are currently many Cre expressing mouse lines that label subsets of neurons, including RGCs (Gong et al., 2007; Ecker et al., 2010; Chen et al., 2011; Drayson et al., 2019; Göz Aytürk et al., 2022). Thus, taking advantage of intersectional approaches to selectively study ipRGCs requires the development of suitable FlpO based mouse models.

Here, we report the development of a new mouse line (*Opn4^FlpO^*) in which FlpO is expressed from the *Opn4* locus. We characterize the number and identity of neurons that express FlpO in the retina, the patterns of axonal projection of these neurons from the retina to multiple downstream targets and confirm that this mouse line is a viable tool for assessing ipRGC-dependent behaviors. Our work provides the field with a new, key resource that will further our understanding of how diverse retinal cell types give rise to light-evoked behavior.

## Material and Methods

### Animal care and use

All experimental procedures related to the use of mice were conducted with approved protocols by the Institutional Animal Care and Use Committee (IACUC) at Northwestern University.

Both male and female mice with a mixed B6/129 mixed background were utilized for all experiments. Mice used for histological examination were between 30 to 90 days of age. Animals used for electrophysiological studies were between 30 to 50 days of age, and those used for behavioral studies were between 60 to 90 days old. The detailed generation of all the animal models is provided below.

### Gene targeting

A bacterial artificial chromosome (BAC) clone containing the *Opn4* gene (bMQ356B1) was isolated from a 129S7/SvEv genomic library (Adams et al., 2005) and transformed into SW102 bacterial cells (Warming et al., 2005). A fragment consisting of an AscI-flanked kanamycin resistance gene was inserted into the BAC via recombineering, replacing a part of *Opn4* exon 1 from the start codon to just after the splice donor (deletion corresponds to GRCm39 chr14:34321773-34321915). A fragment of the modified BAC was retrieved by gap repair into pBluescript with a modified polylinker consisting of XhoI and PmeI sites to generate a targeting vector with a 3.8kb 5’ homology arm and a 4.1kb 3’ homology arm (corresponding to GRCm39 chr14:34325679-34317678). The kanamycin cassette was then excised via AscI digestion and replaced with a fragment comprising the coding sequence for FlpO followed by the rabbit beta globin intron and polyadenylation sequence and a self-excising neomycin selection gene, ACNF (Bunting et al., 1999; Bozza et al., 2002) (Supporting Information Figure S1A).

The targeting vector was linearized with XhoI and electroporated into 129S6 murine embryonic stem cells (PRX129). Neomycin resistant ES clones were selected with G418 and screened by long-range PCR as described (Skarnes et. al., 2011) on 3’ ends using the following primers: P1 (5’-GATATTGCTGAAGAGCTTGGCGGCGAATG-3’) and P2 (5’-GCTGGATCTAGTAATGTTAGGATGGTTGAC-3’) (Supporting Information Figure S1B). Positive clones were injected into C57BL/6J blastocysts, and the resulting chimeras bred to obtain germline transmission of the mutation (Figure 1A and Supporting Information Figure S1A). Germline excision of the ACNF cassette was confirmed via PCR. Opn4^FlpO/FlpO^ mice are fertile and show no overt phenotypes.

**Figure 1.**
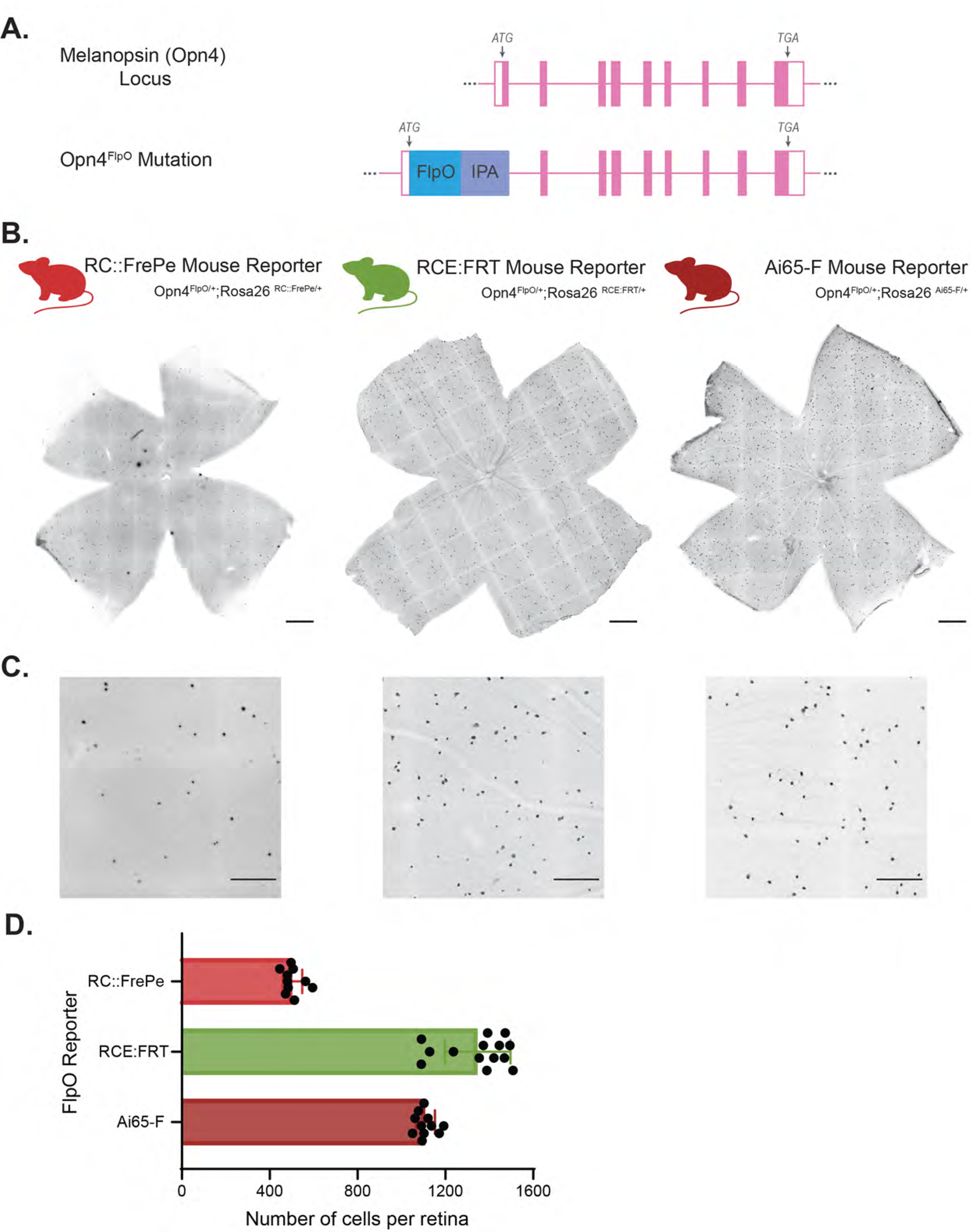
The Opn4^FlpO^ line labels cells in the ganglion cell layer in multiple reporter lines. A. Top, schematic diagram of the mouse melanopsin (Opn4) gene locus. Coding and non-coding exons are shown as filled and unfilled pink boxes, respectively. The location of the start codon (ATG) and the predicted stop codon (TAG) on exon 9 are indicated by the downward facing arrows. For clarity, only one of the known Opn4 splice variants (Pires et al., 2009) is depicted. The coding sequence for FlpO (blue box), followed by an exogenous intron and polyadenylation signal (purple box, IPA) is placed after the start codon replacing exon 1 of melanopsin locus. B. Representative images of whole-mount retinas isolated from mice generated by mating Opn4^FlpO^ animals to multiple FlpO dependent reporter lines. Left, retina from a RC::FrePe (Opn4^FlpO/+^;Rosa26^RC::FrePe/+^) mouse, showed the sparsest cell labeling among the three reporters. Middle, representative retina from a RCE:FRT (Opn4^FlpO/+^;Rosa26^RCE:FRT/+^) animal. Right, retina from an Ai65-F (Opn4^FlpO/+^;Rosa26^Ai65-F/+^) mouse. RCE:FRT and Ai65-F retinas show similar number and distribution of labeled cells. Labeled cells are in the ganglion cell layer. Scale bars 500μm. C. High magnification images from representative retinas in (B). Retinas from different FlpO dependent reporters: left, RC::FrePe, middle, RCE:FRT, and right, Ai65-F. Scale bars 200μm. D. The graph compares the total number of cells per retina from each FlpO dependent. Individual dots on the graph represent the number of labeled cells in one retina. The number of labeled cells per retina was obtained by assessing cell labeling in the ganglion cell layer and the inner nuclear layer. RC::FrePe retinas had an average of 504 ± 45 cells per retina (n=10 retinas from 7 mice). Retinas from RCE:FRT animals had an average of 1347 ± 149 labeled cells per retina (n=14 retinas from 10 mice). Ai65-F retinas had 1109 ± 44 labeled cells per retina (n=11 retinas from 7 mice).

Genomic DNA isolated from tissue was purified and founder mice carrying the FlpO gene were identified by PCR with primer sets P3 (5’-TCACCATCAGACTCTTGTGG-3’) (forward primer in 5’ upstream of the Opn4 locus) and P4 (5’-GCGGCACAGCTGGCGATCTTCTC-3’) (reverse primer in the FlpO coding sequence) to yield a 459 bp product. Founder mice that are heterozygous and carry a wild-type melanopsin allele were identified by PCR with primer sets P4 and P5 (5’-CCTGAAGGAGAGTCCATGCTCA-3’) (reverse primer in exon 1 of the wild type Opn4 gene) to produce a 335 bp amplicon.

The experimental animals used for retinal histology, ex vivo retina recordings, and brain histology were obtained by mating Opn4^FlpO/FlpO^ mice to mice heterozygous for the RC::FrePe, RCE:FRT or Ai65-F transgenes (Bang et al., 2012; Sousa et al., 2009; Daigle et al., 2018) (Supporting Information Figure S2). The offspring from these matings had the following genotypes: Opn4^FlpO/+^;Rosa26^RC::FrePe/+^, Opn4^FlpO/+^;Rosa26^RCE:FRT/+^, and Opn4^FlpO/+^;Rosa26^Ai65-F/+^. All offspring possessed a single allele of the melanopsin (*Opn4*) gene, which is sufficient to retain intrinsic photosensitivity in ipRGCs (Lucas et al., 2003) (Figure 6 and Figure 7). RC::FrePe, RCE:FRT and Ai65-F reporter mouse lines utilizes a CAG (CMV (cytomegalovirus) early enhancer/chicken β-actin) promoter in the ROSA26 locus to drive gene reporter expression (Bang et al., 2012; Sousa et al., 2009; Daigle et al., 2018) (Supporting Information Figure S2A). RC::FrePe is a dual recombinase reporter mouse with a STOP sequence flanked by FRT, Flp recombinase recognition sites, followed by an mCherry sequence and a STOP sequence flanked by loxP, Cre recombinase recognition sites (Bang et al., 2012) (Supporting Information Figure S2A). The presence of FlpO recombinase drives expression of mCherry while expression of GFP is dependent on the presence of both Cre and FlpO recombinases (Bang et al., 2012) (Supporting Information Figure S2A). The single reporter recombinase RCE:FRT harbors neomycin resistance and STOP sequences flanked by FRT sites (Sousa et al., 2009) (Supporting Information Figure S2B). GFP expression reporter expression is driven by the presence of FlpO recombinase (Sousa et al., 2009) (Supporting Information Figure S2B). Ai-65F harbors a STOP sequence flanked by FRT sites (Daigle et al., 2018) (Supporting Information Figure S2C). In this single recombinase reporter, tdTomato fluorescence is driven in cells expressing FlpO recombinase (Daigle et al., 2018) (Supporting Information Figure S2C).

For other experiments, Opn4^FlpO/+^ mice were intercrossed and the resulting Opn4^+/+^, Opn4^FlpO/+^, and Opn4^FlpO/FlpO^ littermates were used for behavioral analyses, and the Opn4^FlpO/+^ offspring were used for intravitreal viral injection and histology.

### Retinal histology

#### Whole-Mount Retinas

To isolate retinas, mice were anesthetized by intraperitoneal (IP) injection of Avertin and sacrificed by cervical dislocation. Eyes were then enucleated and retinas were dissected in 1x PBS. In experiments using retinas from transcardially perfused animals (Supporting Information Figure S3 and Supporting Information Figure S5), eyes were enucleated from perfused animals (described below) and dissected in 1x PBS. Retinas were fixed in 4% paraformaldehyde (Electron Microscopy Sciences) in 1x PBS overnight at 4°C. Retinas were washed with 1x PBS for 1.5 hours (3×30 minutes) at room temperature (RT) and then blocked overnight at 4°C in blocking solution (6% donkey serum in 0.3% Triton PBS). Retinas were then placed in primary antibody solution for 3-4 nights at 4°C. Then, retinas were washed in 1x PBS for 3×30 minutes at RT and incubated in secondary antibody solution for 2 hours at RT for Brn3a labeling, and 4 hours at RT for other antibodies (Supporting Information Figure S5F). Retinas were washed in 1x PBS for 3×30 minutes at RT and mounted using Fluoromount aqueous mounting medium (Sigma).

Primary and secondary antibody solutions were made in 3% normal donkey serum in 0.3% Triton PBS. Primary antibody dilutions were as follows: 1:1000 chicken anti-mCherry (Abcam, Catalog #: ab 205402, RRID:AB_ 2722769), 1:1000 chicken anti-GFP (Abcam, Catalog #: ab13970, RRID:AB_300798), 1:1000 rabbit anti-RBPMS (Millipore, Catalog #: ABN1376, RRID:AB_2687403), 1:2000 rabbit anti-melanopsin (Advanced Targeting Systems, Catalog #: AB-N38, RRID:AB_1608077), 1:1000 mouse anti-SMI-32 (BioLegend, Catalog #: 801701, RRID:AB_509997), and 1:500 mouse anti-Brn3a (Millipore, Catalog #: MAB1585, RRID:AB_94166). Secondary antibody dilutions were as follows: 1:500 donkey anti-chicken Alexa Fluor 488 (Jackson ImmunoResearch Labs, Catalog #: 703-545-155, RRID:AB_2340375), 1:500 donkey anti-rabbit Alexa Fluor 546 (Thermo Fisher Scientific, Catalog #: A10040, RRID:AB_2534016), 1:500 donkey anti-rabbit Alexa Fluor 647 (Thermo Fisher Scientific, Catalog #: A-31573, RRID:AB_2536183), 1:500 donkey anti-mouse Alexa Fluor 546 (Thermo Fisher Scientific, Catalog #: A10036, RRID:AB_11180613), and 1:500 donkey anti-mouse Alexa Fluor 647 (Molecular Probes, Catalog #: A-31571, RRID:AB_162542).

#### Cell Fills

To visualize cells filled during electrophysiological recording, retina pieces were fixed in 4% paraformaldehyde (Electron Microscopy Sciences) in 1x PBS overnight at 4°C. Retinas were washed with 1x PBS for 3×30 minutes at RT and then blocked overnight at 4°C in blocking solution (6% goat serum in 0.3% Triton PBS). Retinas were then placed in primary antibody solution containing 1:1000 mouse anti-SMI-32 (BioLegend, Catalog #: 801701, RRID:AB_509997) in 3% normal donkey serum in 0.3% Triton PBS for 2-4 days at 4°C. Retinas were washed in 1x PBS for 3×30 minutes at RT and transferred to secondary antibody solution containing 1:1000 goat anti-mouse Alexa 488 (Thermo Fisher Scientific, Catalog #: A-21131, RRID:AB_2535771) and 1:1000 streptavidin conjugated with Alexa 546 (Thermo Fisher Scientific, Catalog #: S-11225, RRID:AB_2532130) in 3% normal donkey serum in 0.3% Triton PBS overnight at 4°C. Retinas were then washed in 1x PBS for 3×30 minutes at RT and mounted using Fluoromount aqueous mounting medium (Sigma).

#### Imaging

Whole retina tiled images were captured using a confocal laser scanning microscope (Leica DM5500 SPE, Leica Microsystems) with a 20x objective. To include any sparse labeling in the inner nuclear layer consistent with labeling of ipRGCs, tiled image stacks spanning the ganglion cell layer to inner nuclear layer were collected and automatically assembled (stitched) using Leica Application Suite X (RRID:SCR_013673). Images were processed using Fiji (Schindelin et al., 2012). Dendritic arbors from ipRGCs were traced using the Fiji plugin software, Simple Neurite Tracer (Longair et al., 2011).

### Ex vivo retina preparation for electrophysiology

Mice were dark-adapted overnight and euthanized by CO_2_ asphyxiation followed by cervical dislocation. Eyes were enucleated, and retinas were dissected under dim red light in carbogenated (95% O_2_ - 5% CO_2_) Ames’ medium (Sigma-Aldrich). Retinas were sliced in half and incubated in carbogenated Ames’ medium at 26°C for at least 30 min. Retinas were then mounted on a glass-bottom recording chamber and anchored using a platinum ring with nylon mesh (Warner Instruments). The retina was maintained at 30-32°C and perfused with carbogenated Ames’ medium at a 2-4ml/min flow.

### Solutions for electrophysiology

All recordings were made in Ames’ medium with 23mM sodium bicarbonate. Synaptic transmission was blocked with 100μM DNQX (Tocris), 2μM L-AP4 (Tocris), 100μM picrotoxin (Sigma-Aldrich), and 20μM strychnine (Sigma-Aldrich) in Ames’ medium. 500nM tetrodotoxin (TTX) citrate (Tocris) was added to the synaptic blocker solution for voltage-clamp experiments. The internal solution used contained (in mM): 120 K-gluconate, 5 NaCl, 4 KCl, 10 HEPES, 2 EGTA, 4 ATP-Mg, 0.3 GTP-Na2 and 7 Phosphocreatine-Tris, with the pH adjusted to 7.3 with KOH. The internal solution was passed through a sterile filter with a 0.22 µm pore size (Sigma-Aldrich). Prior to recording, 0.3% Neurobiotin (Vector Laboratories) and 10μM Alexa Fluor 488 (Thermo Fisher) were added to internal solution.

### Electrophysiology and analysis

The ganglion cell layer of retina was visualized using IR-DIC optics at 940 nm. M1, M2, and M3 ipRGCs were targeted in the Opn4^FlpO/+^;Rosa26^Ai65-F/+^ mice line based on their somatic tdTomato signals visualized under brief epifluorescent illumination. Following recording, dendritic stratification was classified by focusing in the proximal and distal layers of the IPL under epifluorescent illumination with an intracellular dye (Alexa 488) and confirmed post-recording by Neurobiotin fill. The identities of M1, M2, and M3 ipRGCs were established using both dendritic stratification, soma size, and light responses. Cells with only OFF ramified dendrites were classified as M1 ipRGCs, cells with only ON stratified dendrites were identified as M2 ipRGCs, and bistratified cells were classified as M3 ipRGCs (M1: OFF; M2: ON; M3 ON/OFF) (Schmidt et al., 2008; Schmidt and Kofuji, 2009; Lee and Schmidt, 2018; Contreras et al., 2023). All cells identified as M2 were confirmed negative for SMI-32 immunolabeling, to ensure any cells with large somata were not M4 ipRGCs (data not shown) (Lee and Schmidt, 2018; Contreras et al., 2023).

The blue LED light (∼480 nm) was used to deliver light stimuli to the retina through a ×60 water-immersion objective. The photon flux was attenuated using neutral density filters (Thor Labs). Before recording, retinas were dark-adapted for at least 5 min. Cells were stimulated with a 50ms full-field flash of bright light with an intensity of 6.08 ×10^15^ photons · cm^−2^ · s^−1^.

Whole-cell recordings were performed using a Multiclamp 700B amplifier (Molecular Devices) and fire-polished borosilicate pipettes (Sutter Instruments, 3-6 MΩ). All voltage traces were sampled at 10 kHz, low-pass filtered at 2 kHz, and acquired using a Digidata 1550B and pClamp 10 software (Molecular Devices). All reported voltages are corrected for a −13 mV liquid junction potential calculated using Liquid Junction Potential Calculator in pClamp.

Electrophysiology recordings were analyzed using custom scripts written in MATLAB (MathWorks; RRID:SCR_001622). For experiments measuring the intrinsic melanopsin response to a 50ms light stimulus (6.08 × 10^15^ photons · cm^−2^ · s^−1^), cells were voltage clamped at −60 mV. We measured the maximum amplitude of the light response for each cell, using a similar analysis described in Contreras et al., 2023. Cells were smoothed using a 100ms sliding average. The baseline current was quantified as the mean current in the first 3 s of the recording protocol. Maximum amplitude was calculated as the maximum current value of the smoothed trace that had the greatest change from baseline during the recording period. Values were reported as the absolute value of the current (pA).

### Viral intravitreal Injections

Mice between P50-60 were anesthetized by intraperitoneal (IP) injection of Avertin (2,2,2-Tribromoethanol) and a 30-gauge needle was used to open a hole in the ora serrata. Each eye was injected with 1μL of AAV2-Ef1a-fDIO-mCherry (7×10¹² vg/mL; Addgene, Catalog #: 114471-AAV2) using a custom Hamilton syringe (Borghuis Instruments) with a 33-gauge needle (Hamilton). Mice were monitored until they recovered from anesthesia.

### Brain histology

Mice were anesthetized by IP injection of Avertin and transcardially perfused with PBS followed by 4% paraformaldehyde (Electron Microscopy Sciences) in PBS. Brains were harvested and post-fixed overnight in 4% PFA. Brains were washed with PBS three times and cryoprotected in a 30% sucrose solution in PBS for two nights at 4°C. Brains were then embedded and frozen in OCT using dry ice. 100µm coronal sections were made on a Leica CM1950 cryostat. Sections were collected in PBS and blocked in 6% normal goat serum in 0.3% Triton PBS overnight at 4°C. Sections were then transferred to primary antibody solution containing rabbit anti-dsRed (1:1000, Takara, Catalog #: 632496, RRID:AB_10013483) in 0.3% Triton PBS with 3% normal goat serum for 2 nights at 4°C. Sections were washed in PBS for 3×30 minutes at room temperature (RT) and transferred to secondary antibody solution overnight at 4°C. Secondary antibody solution contained Alexa 546 goat anti-rabbit (1:1000, Thermo, Catalog #: A-11035, RRID:AB_143051) in 3% normal goat serum in 0.3% Triton PBS. Sections were then washed in PBS at RT and mounted on glass cover slides with Vectashield mounting medium (Vector Laboratories) and imaged using a confocal laser scanning microscope (Leica DM5500 SPE, Leica Microsystems).

### Antibody Characterization

The primary antibodies used in this study are listed in Table 1. These primary antibodies have been characterized in the mammalian retina and brain; additionally, our staining patterns match those documented in other studies. *<u>mCherry antibody:</u>* According to the manufacture the mCherry antibody was raised against the full-length mCherry protein and recognizes both tdTomato and mCherry proteins (Shaner et al., 2004; Sun et al., 2022). The mCherry antibody staining patterns have been reported in other ipRGC studies (Sonoda et al., 2020a; Beier et al., 2021; Berry et al., 2023). *<u>GFP antibody:</u>* The GFP (Green Fluorescent Protein), antibody was raised against full-length GFP protein. Its specificity has been demonstrated in a variety of systems including transfected cells and the mammalian retina (Beier et al., 2021; Zhang, et al., 2021). *<u>RBPMS</u> <u>antibody:</u>* RBPMS (RNA-binding protein with multiple splicing), has one RRM (RNA recognition motif) domain and belongs to the RRM family of RNA-binding proteins. RBPMS is considered a pan-RGC marker, and reports have shown that this antibody exclusively labels RGCs in mammalian species (Rodriguez et al., 2014; Beier et al., 2021). *<u>Melanopsin antibody:</u>* The melanopsin antibody was raised against a synthetic polypeptide consisting of the 15 N-terminal amino acids of mouse melanopsin with an additional C-terminal cysteine (MDSPSGPRVLSSLTQC) conjugated to KLH (Provencio et al., 2002). The specificity of this antibody has been confirmed in studies demonstrating a lack of immunoreactivity in the retinas from Opn4^−/−^ mice (Panda et al., 2002). This antibody immunolabels M1, M2, and M3 ipRGC subtypes (Berson et al., 2010). *<u>SMI-32 antibody:</u>* The SMI-32 antibody was generated against a non-phosphorylated epitopes of on the medium and heavy molecular weight subunits of neurofilament H (NFH) (Sternberger and Sternberger, 1983). This antibody labels cell bodies dendrites, and thick axons (Sternberger and Sternberger, 1983). SMI-32 is a marker of ON-sustain alpha RGCs which are synonymous with M4 ipRGCs (Coombs et al., 2006; Bleckert et al., 2014; Schmidt et al., 2014; Krieger et al., 2017; Sonoda et al., 2018; Sonoda et al., 2020b). *<u>Brn3a antibody:</u>* The Brn3a antibody was raised against the POU-domain (amino acids 186µ224) of Brn3a fused to the T7 gene 10 protein. According to the manufacture, this antibody only recognizes Brn3a but not Brn3b or Brn3c. The Brn3a antibody has been shown to label broad RGC populations except for ipRGCs (Quina et al., 2005; McNeill et al., 2011). *<u>ds-Red antibody:</u>* According to the manufacture, this antibody was generated against DsRed-Express, a variant of *Discosoma sp.* Red fluorescent protein. The ds-Red antibody recognizes the ds-Red, mCherry and tdTomato. The specificity of this antibody has been confirmed in Figure S9. Brain sections from Opn4^−/−^ mice showed no ds-Red immunoreactivity (Figure S9).

**Table 1.**
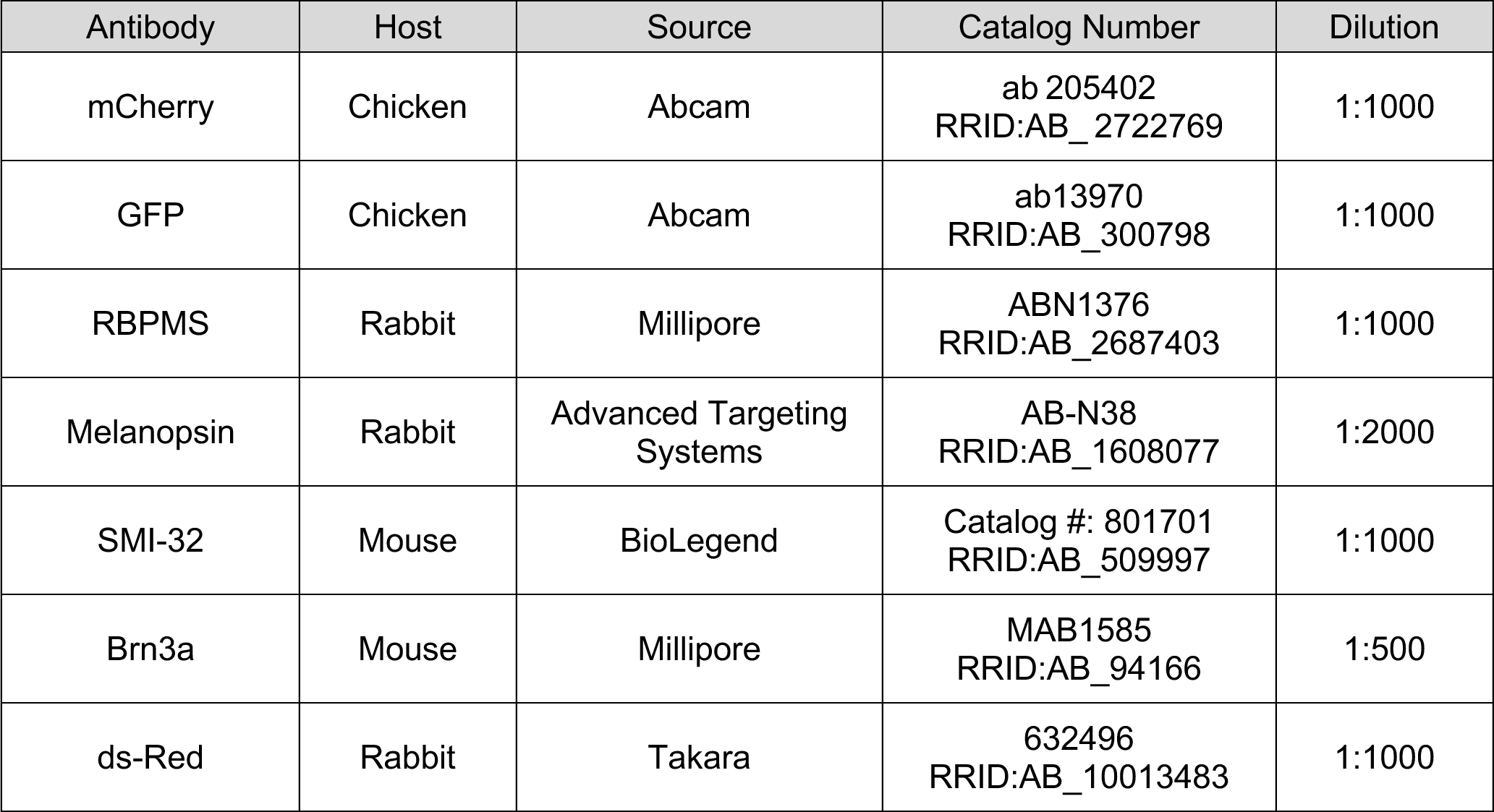
Primary Antibodies.

### Pupillometry

Consensual pupillary light reflex (PLR) and post-illumination pupil recovery (PIPR) testing was measured in male and female mice by a researcher blinded to genotype groups. Mice were acclimated to handling and mock PLR testing for at least 4 days before data was collected. Tests were performed in dim red light between ZT8 and ZT12, after at least 1 hour of dark adaptation. For each test, the mouse was securely scruffed and held on a platform on top of a clean paper towel, with the left eye positioned ∼45 mm from a 480 nm Schott KL 1600 LED light source, fitted with neutral density filters. The recording area was illuminated by infrared light. 60 fps recordings of the contralateral (right) eye were captured in OBS Studio with a DMK 22AUC03 USB camera (The Imaging Source) with an infrared bandpass filter and 1/2.5” 4.0 mm lens with manual focus. The pupil was recorded for ∼5 seconds in the dark before the light stimulus was switched on for 20 seconds to capture the PLR, and the recording continued in the dark for 25 seconds to capture the PIPR. This was repeated at moderate (13.9 log photons· cm^−2^ · s^−1^) and bright light levels (14.9 log photons· cm^−2^ · s^−1^), on separate days of testing. As autonomic tone influences PLR, mice that displayed anxiety-like behaviors (struggling/vocalizing, accompanying obvious changes in pupil size) were re-tested on a new day. Recordings from 1-2 mice were excluded from each light level due to persistent anxiety-like responses.

PLR and PIPR were quantified using a machine learning model trained for 350k iterations in DeepLabCut (Mathis et al., 2018) using 180 labeled frames from 18 videos, and validated against results from 3 random subjects quantified manually in Fiji.(Schindelin et al., 2012) (Supporting Information Figure S7) For each frame, the area of the pupil as a ratio of the eye area were calculated from DeepLabCut-generated XY coordinates. These data were refined in two passes, first by manual removal of “impossible” data points generated by incorrect tracking, and then by removing any data points falling more than 2 standard deviations from the mean of each set of 12 frames (0.2 seconds). Data were then normalized to the size of the dark pupil and analyzed at the 0.2 second resolution. The dark pupil size was calculated as the average pupil size during the last 2 seconds of recording before the light stimulus, PLR was analyzed from the first frame of the light stimulus for 20 seconds, and PIPR from the first frame of post-stimulus darkness for 25 seconds. For PLR and PIPR, dark pupil size, maximum constriction, recovery, and time constants (Tau) were compared using two-way ANOVA or mixed effects analysis with Tukey post-hoc testing. Group results over time were compared using repeated measures mixed models. Kinetics were compared by fitting single phase exponential decay functions to the 0.2 second resolution datasets for each mouse to determine individual Tau values.

## Results

### Generation of *Opn4^FlpO^* mouse line

To allow for cell-type specific manipulation of ipRGCs without the use of Cre recombinase, we developed a strain of mice (Opn4^FlpO^) in which the recombinase FlpO is expressed under the control of the endogenous Opn4 regulatory sequences. To do this, we used gene targeting in embryonic stem (ES) cells to insert the FlpO coding sequence at the Opn4 start codon in exon 1, followed by an exogenous intron and polyadenylation signal (Supporting Information Figure S1A). The resulting allele disrupts the Opn4 coding sequence and is predicted to terminate the endogenous Opn4 transcript due to the presence of the polyadenylation sequence therefore producing a loss of Opn4 function (Figure 1A and Supporting Information Figure S1A).

### Number of retinal neurons labeled in Opn4^FlpO^ mouse line

We first sought to determine the pattern of recombinase activity in our novel Opn4^FlpO^ line. Previous work with a different Opn4 driver (Opn4^Cre^) showed that the number of neurons showing Cre-dependent labeling varied with the reporter gene used. Crosses with Z/AP or Z/EG reporters labeled 3-fold fewer RGCs than crosses with more sensitive, Rosa-based Cre reporters (Ecker et al., 2010; Maloney et al., 2024). Because reporter lines can vary in the number of cells labeled and the fidelity of reporter expression, we characterized the total number, distribution, and location of retinal neurons labeled when Opn4^FlpO^ animals were crossed to multiple FlpO dependent reporter lines: RC::FrePe, RCE:FRT, and Ai65-F. All three reporters contain a CAG promoter in the ROSA26 locus to drive reporter gene expression (Supporting Information Figure S2) (Sousa et al., 2009; Bang et al., 2012; Daigle et al., 2018). RC::FrePe, is a dual recombinase mouse reporter in which expression of mCherry is dependent on the presence of FlpO recombinase alone and expression of GFP is dependent on the presence of both Cre and FlpO recombinases (Bang et al., 2012) (Supporting Information Figure S2A). RCE:FRT and Ai65-F, on the other hand, are single-recombinase reporters with FlpO-dependent GFP or tdTomato expression respectively (Supporting Information Figure S2A-B) (Sousa et al., 2009; Daigle et al., 2018). All crosses examined were compound heterozygous for the driver and reporter alleles. Crosses with all three reporters primarily labeled cells in the ganglion cell layer, along with sparse labeling in the inner nuclear layer, consistent with labeling of displaced ipRGCs. The RCE:FRT and Ai65-F reporters showed similar labeling efficiencies, with an average of 1347 ± 149 labeled cells per retina (n=14 retinas from 10 mice) and 1109 ± 44 labeled cells per retina (n=11 retinas from 7 mice) respectively (Figure 1B-D). The RC::FrePe reporter showed the sparsest cell labeling, with an average of 504 ± 45 cells per retina (n=10 retinas from 7 mice) (Figure 1B-D). This sparse labeling in RCE::FrePe animals has been previously observed in other neurons (Plummer et al., 2015). Notably, all lines labeled fewer cells than reported in Opn4^Cre^, which typically labels over 2000 cells per retina when crossed to both low- and high-efficiency reporters. (Ecker et al., 2010). This difference is likely due to the lower efficiency of FlpO recombinases (Ecker et al., 2010; Maloney et al., 2024; Andreas et al., 2002; Schaft et al., 2001; Kranz et al., 2010; Zhao et al., 2023). Overall, we find that the RCE-FRT is the most sensitive reporter of Opn4^FlpO^ expression in fresh fixed tissue.

To examine whether tissue fixation influenced the efficiency of labeling, we also quantified cell numbers in retinas from animals that were fixed via perfusion rather than via immersion fixation. Under these conditions, the number of cells labeled in Ai65-F retinas doubled to 2117 ± 101 cells per retina (n=8 retinas from 4 mice), comparable, though slightly lower, than the number of cells seen when Opn4^Cre^ is crossed to insensitive, but high fidelity, Z/AP and Z/EG Cre dependent reporters (Supporting Information Figure S3) (Ecker et al., 2010; Maloney et al., 2024). Notably, this Opn4^FlpO^ line labels significantly fewer cells when compared with the Opn4^Cre^ line crossed to Ai lines, a combination that results in extensive ectopic labeling of other RGCs (Maloney et al., 2024). This suggests that the labeling seen by the Opn4^FlpO^ line crossed to higher sensitivity reporters and under optimized fixation conditions is not the result of ectopic labeling.

### Identity of neurons labeled in Opn4^FlpO^ mouse line

The FlpO-recombinase in the Opn4^FlpO^ line should be selectively expressed in ipRGCs and not other types of RGCs or retinal neurons. To test this, we used immunohistochemistry to determine the extent of overlap between labeling in all three reporter lines and four markers of various retinal cell populations: RBPMS (a pan-RGC marker), Opn4 (a marker of M1-M3 ipRGCs), SMI-32 (a marker of alpha RGCs, including M4 ipRGCs), and Brn3a (an RGC-specific transcription factor that is absent from ipRGCs) (Rodriguez et al., 2014; Provencio et al., 2002; Ecker et al, 2010; Berson et al., 2010; Quattrochi et al., 2019; Schmidt et al., 2014; Lee and Schmidt, 2018; Sonoda et al., 2018; Sonoda et al., 2020b; Quina et al., 2005; McNeill et al., 2011).

We first assessed whether the cells labeled in Opn4^FlpO^ animals are RGCs. If this is the case, then all fluorescently labeled cells in a given reporter line should co-label with the pan-RGC marker RBPMS. Indeed, for RC::FrePe (n=4), RCE:FRT (n=3), and Ai65-F (n=5) retinas, greater than 99% of cells expressing the fluorescent reporter co-labeled with RBPMS (Figure 2).

**Figure 2.**
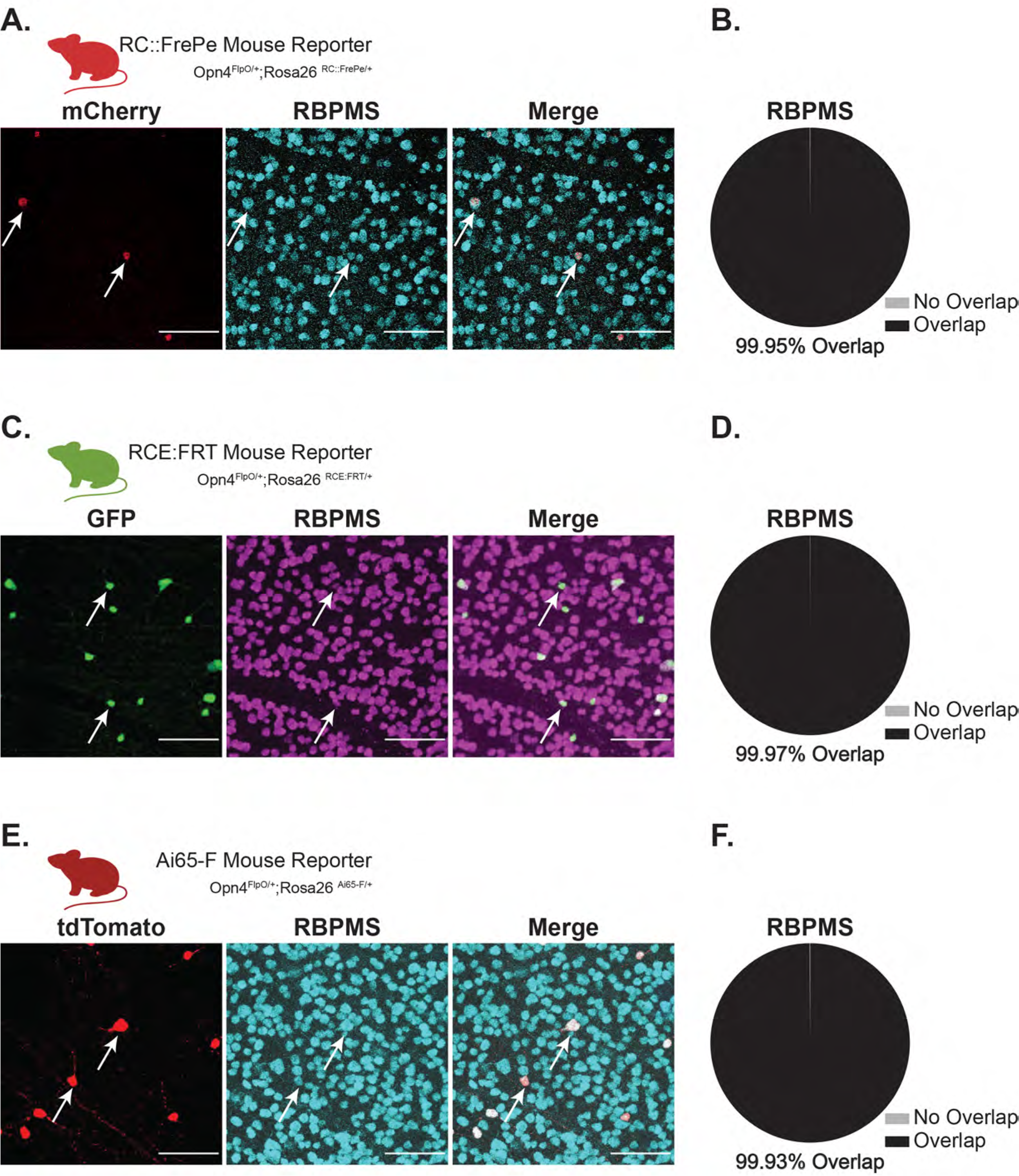
Cells labeled in the Opn4^FlpO^ mouse line are retinal ganglion cells (RGCs). A. Representative images of the ganglion cell layer in a whole-mount retina from a RC::FrePe (Opn4^FlpO/+^;Rosa26^RC::FrePe/+^) animal immunolabeled for mCherry (red, left panel) and the pan-RGC marker RBPMS (cyan, middle panel). Right panel shows the merged image. B. Total proportion of mCherry cells co-labeled (black) or not (gray) with RBPMS in RC::FrePe retinas (n=4 retinas from 3 mice). C. Representative images of the ganglion cell layer in a whole-mount retina from a RCE:FRT (Opn4^FlpO/+^;Rosa26^RCE:FRT/+^) mouse immunolabeled for GFP (green, left panel) and RBPMS (purple, middle panel). Right panel shows merged image. D. Total proportion of GFP cells co-labeled (black) or not (gray) with RBPMS in RCE:FRT retinas (n=3 retinas from 2 mice). E. Representative images of the ganglion cell layer in a whole-mount retina from a Ai65-F (Opn4^FlpO/+^;Rosa26^Ai65-F/+^) mouse immunolabeled for tdTomato (red, left panel) and RBPMS (cyan, middle panel). Right panel shows merged image. F. Total proportion of tdTomato cells co-labeled (black) or not (gray) with RBPMS in Ai65-F retinas (n=5 retinas from 3 mice). White arrows show example reporter cells co-labeled with RPBMS. Scale bar 100μm.

While all ipRGCs express Opn4, the levels of expression vary systematically by subtype. M1-M3 ipRGCs express relatively higher levels of Opn4 than M4-M6. To determine which subpopulations of ipRGCs are marked, we stained tissue using an Opn4 antibody, which is known to label M1-M3 ipRGCs. We observed that 83.99% of mCherry+ cells in RC::FrePe retinas (n = 3), 89.53% of GFP+ cells in RCE:FRT retinas (n=5), and 69% of tdTomato+ cells in Ai65-F retinas (n = 2) are Opn4 positive in each reporter (Figure 3). These results suggest that the majority of cells labeled in the RC::FrePE line are M1-M3 ipRGCs, while additional cell types are also labeled in the RCE:FRT and Ai65-F lines.

**Figure 3.**
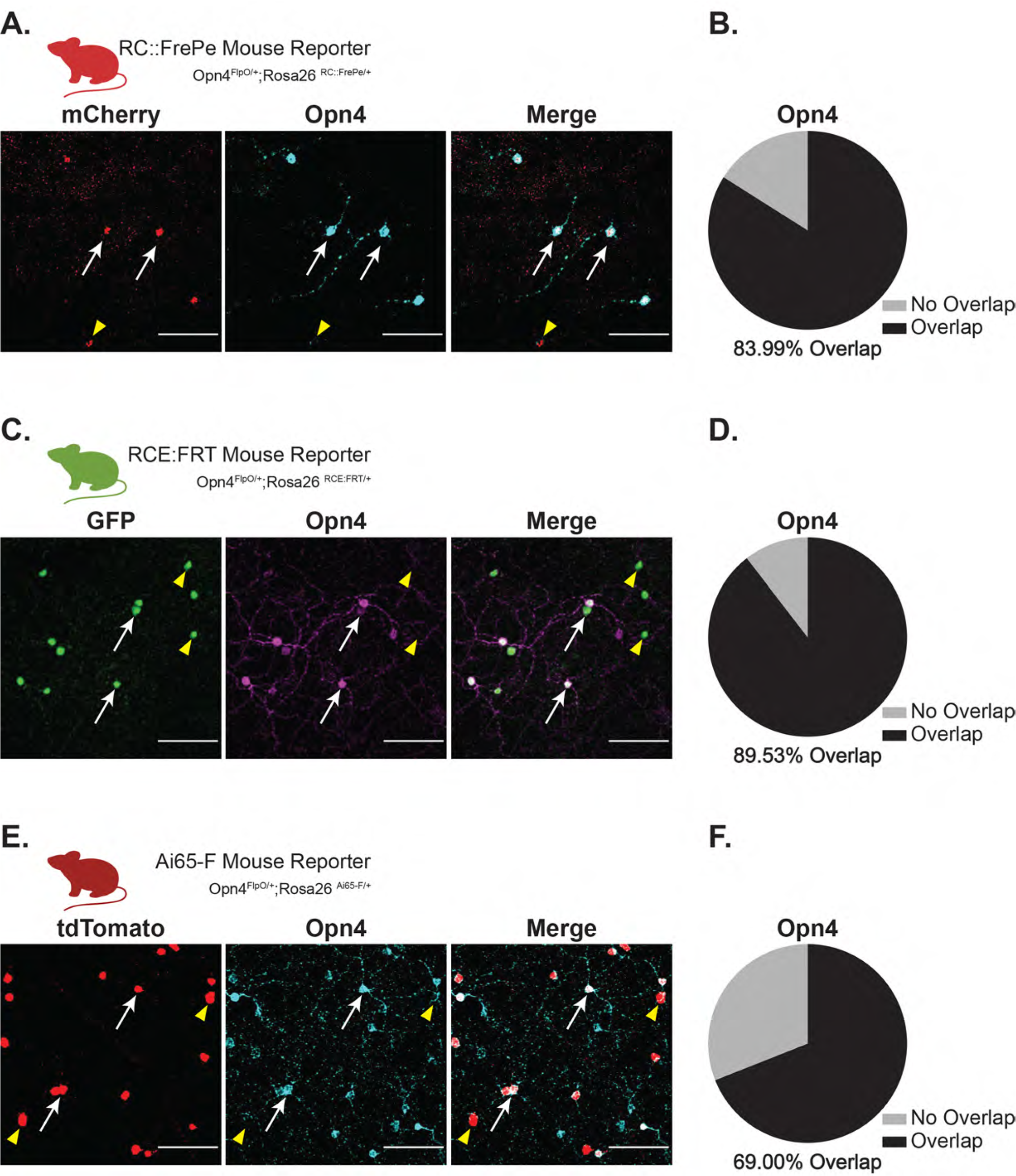
The majority of labeled cells in the Opn4^FlpO^ mouse line Opn4 immunoppositive. A. Representative images of the ganglion cell layer in a whole-mount retina from a RC::FrePe (Opn4^FlpO/+^;Rosa26^RC::FrePe/+^) animal immunolabeled for mCherry (red, left panel) and Opn4 (cyan, middle panel). Right panel shows merged image. B. Total proportion of mCherry cells co-labeled (black) or not (gray) with in RC::FrePe retinas (n=3 retinas from 2 mice). C. Representative images of the ganglion cell layer in a whole-mount retina from a RCE:FRT (Opn4^FlpO/+^;Rosa26^RCE:FRT/+^) mouse immunolabeled for GFP (green, left panel) and Opn4 (purple, middle panel). Right panel shows merged image. D. Total proportion of GFP cells co-labeled (black) or not (gray) with Opn4 in RCE:FRT retinas (n=5 retinas from 3 mice). E. Representative images of the ganglion cell layer in a whole-mount retina from a Ai65-F (Opn4^FlpO/+^;Rosa26^Ai65-F/+^) mouse immunolabeled for tdTomato (red, left panel) and Opn4 (cyan, middle panel). Right panel shows merged image. F. Total proportion of tdTomato cells co-labeled (black) or not (gray) for Opn4 in Ai65-F retinas (n=2 retinas from 2 mice). White arrows show example reporter cells co-labeled with Opn4. Yellow triangles illustrate reporter cells without any Opn4 overlap. Scale bar 100μm.

We next labeled retinas for SMI-32, which labels all M4 ipRGCs as well as other alpha RGC types. In the RC::FrePe line, we observed that 0.60% (n =2) of mCherry cells were labeled with SMI-32 (Figure 4A-B). Slightly higher percentages of overlap were seen in RCE:FRT (5.58%; n = 4; Figure 3F-G) and in Ai65-F (12.32%; n = 4) retinas (Figure 4C-F). This indicates that some M4 ipRGCs are likely labeled in the RCE:FRT and Ai65-F lines. The reason for the difference in ratios of M1-M3 versus M4 ipRGCs between the RCE:FRT and Ai65-F lines is unclear, and potentially due to different recombination efficiencies of these constructs across different cell types. The RCE:FRT line is best for avoiding expression in M4 cells. Regardless, this variance highlights the differences in efficiencies that exist across different reporters.

**Figure 4.**
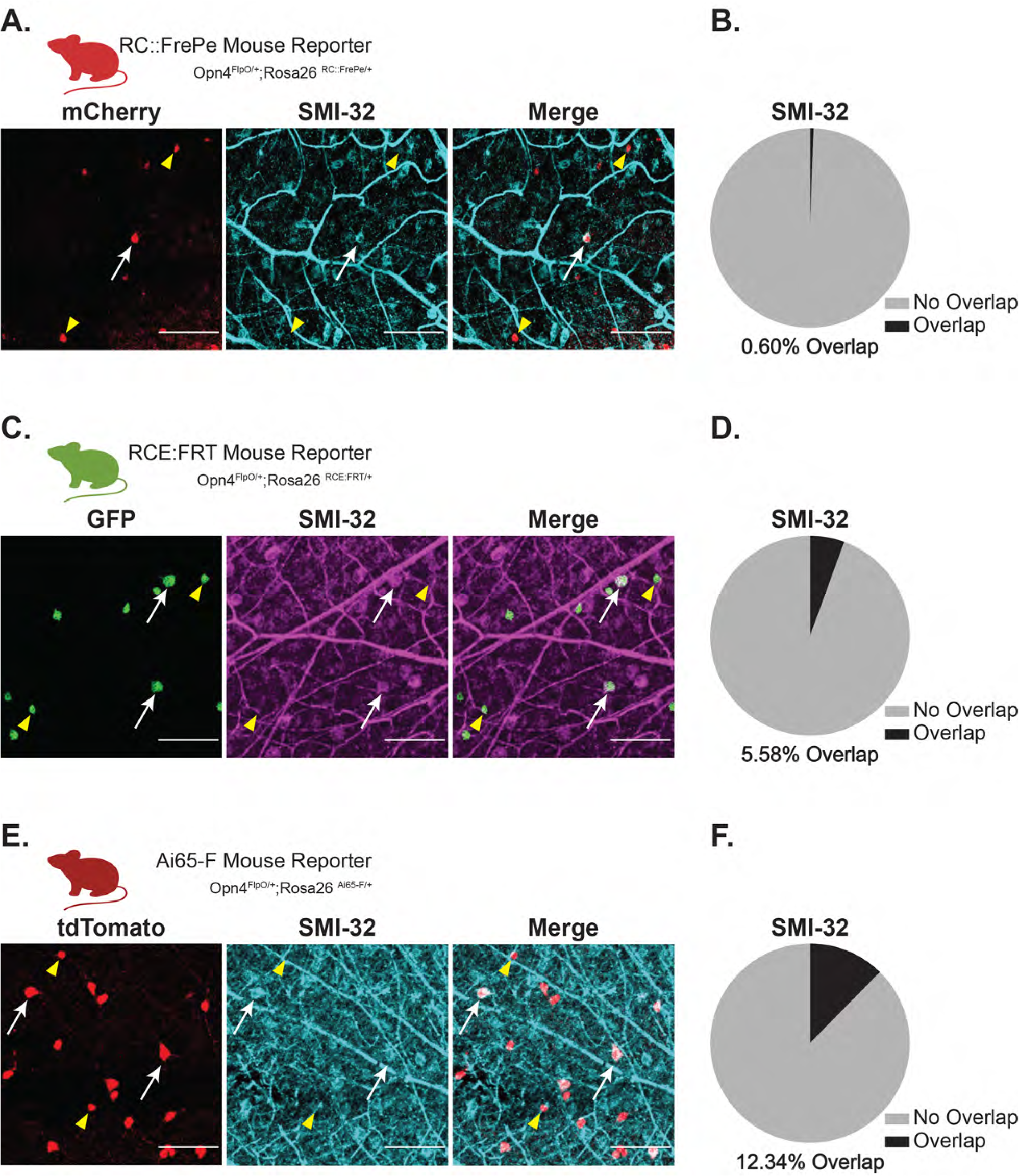
Opn4^FlpO^ mouse line labels some SMI-32-positive retinal ganglion cells. A. Representative images of the ganglion cell layer in a whole-mount retina from a RC::FrePe (Opn4^FlpO/+^;Rosa26^RC::FrePe/+^) animal immunolabeled for mCherry (red, left panel) and SMI-32 a marker of alpha RGCs, including M4 ipRGCs (cyan, middle panel). Right panel shows merged image. B. Total proportion of mCherry cells co-labeled (black) or not not (gray) with SMI-32 in RC::FrePe retinas (n=2 retinas from 2 mice). C. Representative images of the ganglion cell layer in a whole-mount retina from a RCE:FRT (Opn4^FlpO/+^;Rosa26 ^RCE:FRT/+^) mouse immunolabeled for GFP (green, left panel) and SMI-32 (purple, middle panel). Right panel shows merged image. D. Total proportion of GFP cells co-labeled (black) or not (gray) with SMI-32 in RCE:FRT retinas (n=4 retinas from 3 mice). E. Representative images of the ganglion cell layer in a whole-mount retina from a Ai65-F (Opn4^FlpO/+^;Rosa26^Ai65-F/+^) mouse immunolabeled for tdTomato (red, left panel) and SMI-32 (cyan, middle panel). Right panel shows merged image. F. Total proportion of tdTomato positive cells co-labeled (black) or not (gray) with SMI-32 in Ai65-F retinas (n=4 retinas from 2 mice). White arrows denote reporter cells co-expressing SMI-32 and yellow triangles illustrate reporter cells with no SMI-32 overlap. Scale bar 100μm.

In addition to the overlap with ipRGC markers, we also sought to confirm that Opn4^FlpO^ mice are not driving reporter expression in non-ipRGCs. To test this, we examined the overlap of fluorescent reporter expression with labeling for Brn3a, which is not expressed by ipRGCs but is expressed in numerous other RGC types (Quina et al., 2005; McNeill et al., 2011). Across all three reporter lines, we observed that well below 1% of fluorescent cells in all FlpO reporters were labeled with Brn3a (Figures 5). This indicates that the Opn4^FlpO^ allele does not drive widespread reporter expression in ectopic cell types. The proportion of labeling for Opn4, RBPMS, and Brn3a was remarkably similar across reporters (Supporting Information Figure S4). The increase in SMI-32 is likely driven by the higher recombination in RCE:FRT and Ai65-F reporter lines, and is supported by the increased number of cells labeled in these lines and the decrease in proportion of Opn4 cells labeled in Ai65-F retinas, which had the highest proportion of SMI-32 overlap. Indeed, even perfused Ai65-F retinas, which have double the number of tdTomato+ neurons, show highly similar proportions of RBPMS, SMI-32, and Brn3a labeling as the immersion fixed Ai65-F retinas (Supporting Information Figure S5 and Supporting Information Figure S6). Collectively, these results indicate that the Opn4^FlpO^ line drives FlpO-dependent reporter expression primarily in M1-M3 ipRGCs, as well as a small proportion of M4 ipRGCs. Likewise, the lack of Brn3a expression in any of the three lines suggests that, despite some variation in the proportion of ipRGC subtype labeled in different reporter lines, the majority of cells labeled in all three are ipRGCs.

**Figure 5.**
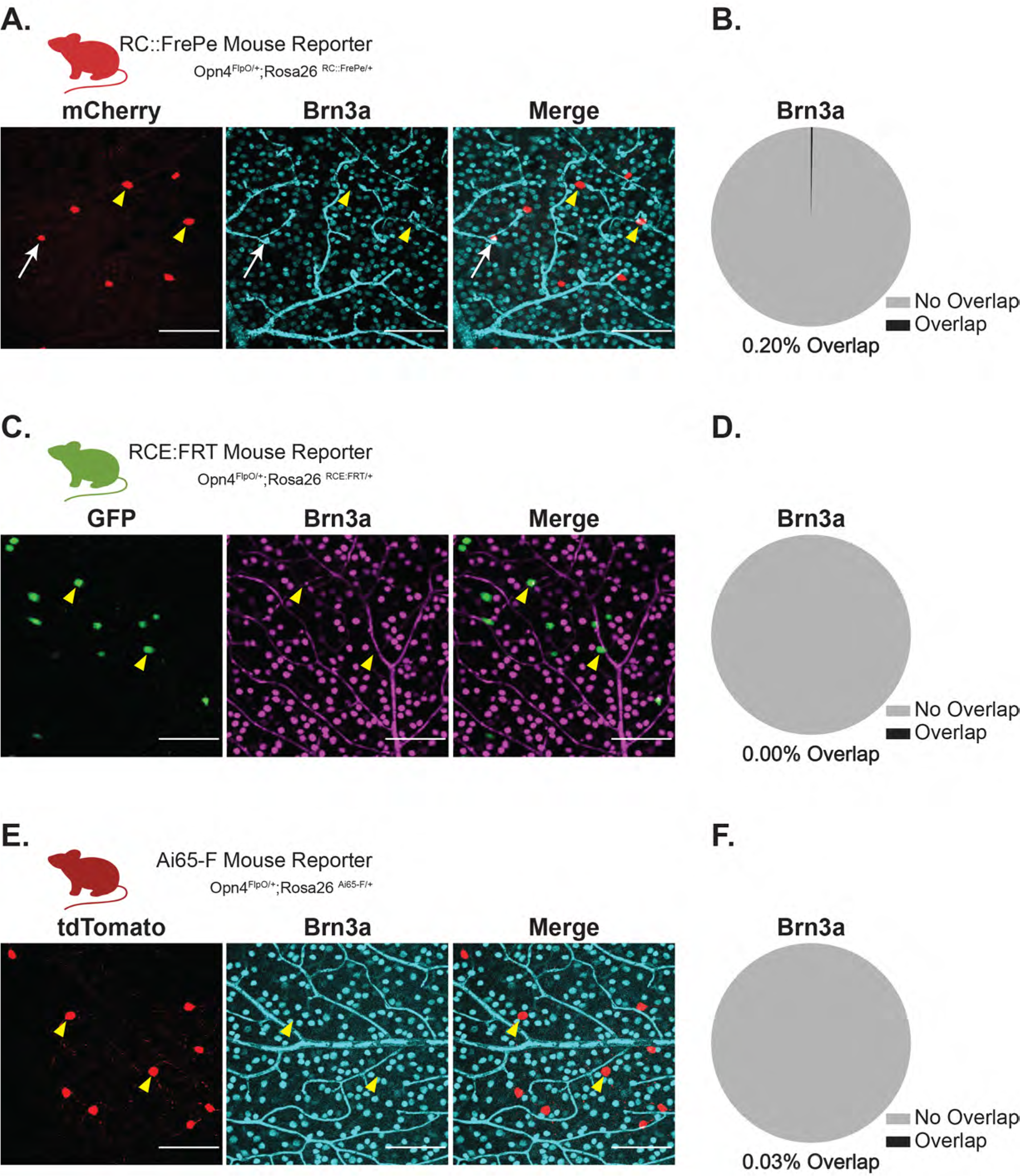
Opn4^FlpO^ mouse line does not label Brn3a-positive conventional retinal ganglion cells. A. Representative images of the ganglion cell layer in a whole-mount retina from a RC::FrePe (Opn4^FlpO/+^;Rosa26^RC::FrePe/+^) animal immunolabeled for mCherry (red, left panel) and conventional RGC marker Brn3a (cyan, middle panel). Right panel shows merged image. B. Total proportion of mCherry cells co-expressing (black) or not (gray) Brn3a in RC::FrePe retinas. Only 0.20% of the total mCherry cells co-labeled with Brn3a in RC::FrePe retinas (n=2 retinas from 2 mice). C. Representative images of the ganglion cell layer in a whole-mount retina from a RCE:FRT (Opn4^FlpO/+^;Rosa26 ^RCE:FRT/+^) mouse immunolabeled for GFP (green, left panel) and Brn3a (purple, middle panel). Right panel shows merged image. D. Total proportion of GFP cells co-labeled (black) or not (gray) with Brn3a in RCE:FRT retinas. No GFP cells co-expressed Brn3a in RCE:FRT retinas (n=3 retinas from 2 mice). E. Representative images of the GCL in a whole-mount retina from a Ai65-F (Opn4^FlpO/+^;Rosa26^Ai65-F/+^) mouse immunolabeled for tdTomato (red, left panel) and Brn3a (cyan, middle panel). Right panel shows merged image. F. Total proportion of tdTomato positive cells co-labeled (black) or not (gray) with Brn3a in Ai65-F retinas (n=3 retinas from 2 mice). White arrow shows an example cell co-expressing Brn3a and fluorescent reporter, and yellow triangles denote reporter-positive cells lacking Brn3a labeling. Scale bar 100μm.

### RGCs in Ai65-F retinas are intrinsically photosensitive

In order to confirm the identity of labeled cells as ipRGCs, and to test their viability, we measured their intrinsic photosensitivity, morphology, and electrophysiological properties using patch clamp recording. An advantage of this approach is that it allows for definitive identification of ipRGC subtypes based on well-established morphological and physiological criteria (see Methods). Of the two most sensitive reporters (RCE:FRT and Ai65-F), the Ai65-F retinas showed readily visible tdTomato under epifluorescent illumination while GFP fluorescence was dim and difficult to detect in ex vivo retinas. Thus, we performed targeted whole-cell patch clamp recordings of tdTomato positive neurons in ex vivo retinas from Opn4^FlpO^/Ai65-F mice. Recordings were made under pharmacological blockade to isolate the intrinsic, melanopsin-based light response. We recorded from cells in voltage clamp mode at a holding potential of −60mV and stimulated cells with a 50 ms, full-field flash of bright 480 nm (6.08×10^15^ photons· cm^−2^ · s^−1^) light. All cells were filled with Neurobiotin and post-fixed for subtype identification post-recording.

Using this approach, we recorded from a total of 15 cells, all of which were intrinsically photosensitive These cells were all identified as M1, M2, or M3 cells (see Methods). The morphological features and photocurrent amplitudes matched those reported previously for all three subtypes (Figure 6) (Baver et al., 2008; Berson et al., 2002, 2010; Hattar et al., 2002, 2006; Viney et al., 2007; Schmidt and Kofuji, 2009, 2011; Müller et al., 2010; Ecker et al., 2010; Estevez et al., 2012; Schmidt and Kofuji, 2011). We did observe a few putative M4 cells that were tdTomato+ in ex vivo Ai65-F retinas that had very large somata, but we were unable to successfully record from these to confirm their identity. Collectively, these results confirm that our FlpO driver preferentially labels M1-M3 ipRGCs and that ipRGC morphology and cellular function are intact. Thus, the Opn4^FlpO^ mouse line is a viable model for ipRGC manipulation without disruption of cellular structure or function.

**Figure 6.**
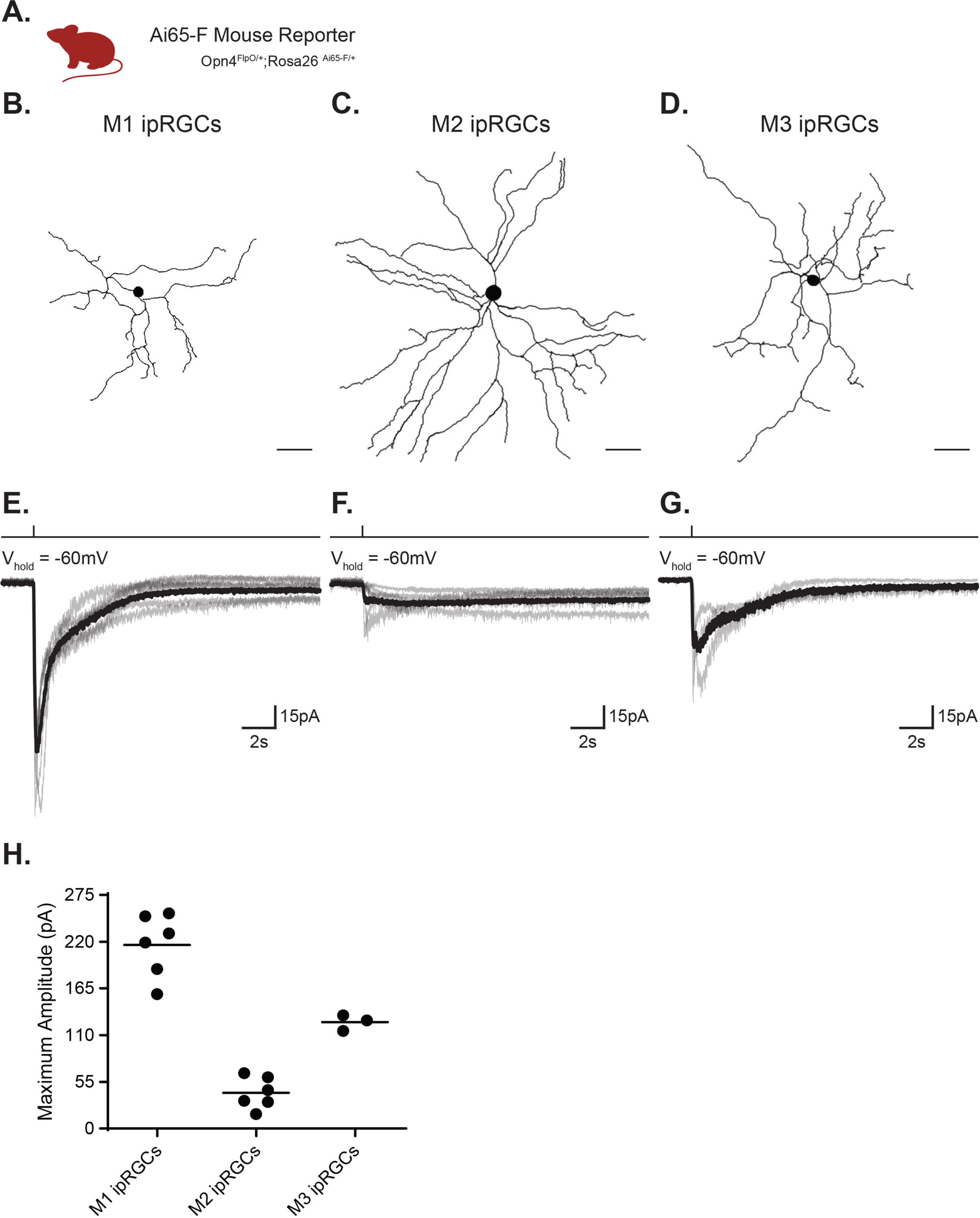
Cells labeled in Ai65-F mouse line are intrinsically photosensitive. A. Whole-cell voltage-clamp recordings and post-hoc cell fills were performed in retinas from Ai65-F (Opn4^FlpO/+^;Rosa26^Ai65-F/+^) animals. B. – C. Representative cell tracing of filled M1, M2, and M3 ipRGCs recorded via whole cell patch clamp. Scale bar 50μm. E. – G. Whole-cell voltage-clamp recording of M1, M2, and M3 ipRGC photocurrents. Individual light responses of M1 (n = 6), M2 (n=6), M2 (n=3) are shown in gray and corresponding average trace is overlayed in black. H. Absolute value of the current amplitudes for the light responses in (E. – G.). Cells were stimulated with 50ms, full-field 480nm light (6.08 × 10^15^ photons · cm^−2^ · s^−1^) pulse of bright light in the presence of synaptic blockers to isolate the intrinsic melanopsin response.

### Opn4^FlpO/FlpO^ animals show pupil constriction deficits similar to other Opn4^−/−^ lines

The Opn4^FlpO^ allele is predicted to produce a loss-of-function of Opn4. Previously generated Opn4^−/−^ mice show deficits in consensual (contralateral) pupil constriction under very bright, 480nm light (Güler et al., 2008; Ecker et al., 2008; Hatori et al., 2008; Le et al., 2008; Xue et al., 2011; Do et al. 2011; Do, 2019). To test whether Opn4^FlpO/FlpO^ animals show similar pupillary response deficits, we measured the consensual pupillary light response (PLR) in Opn4^FlpO/FlpO^, Opn4^FlpO/+^, and Opn4^+/+^ littermates (n = 6-7 per group, Figure 7A) to 480 nm light delivered at bright (14.9 log photons· cm^−2^ · s^−1^) and moderate (13.9 log photons· cm^−2^ · s^−1^) light intensities, following 1 hour of dark adaptation (Figure 7B,G,I). Pupil diameter was measured with DeepLabCut (Mathis et al., 2018 and see Methods), which yielded rapid measurements of pupil diameter that were similar to those obtained using manual methods (Supporting Information Figure S7). Resting pupil diameter in the dark was similar across genotypes (Supporting Information Figure S7). In contrast, Opn4^FlpO/FlpO^ mice showed significantly less constriction than Opn4^FlpO/+^ and Opn4^+/+^ animals in response to light (Figure 7C). We quantified the tau of the response kinetics by fitting one-phase exponential decay functions to the animals from each genotype and condition and found that these were similar for all genotypes at both light levels (Figure 7D). Opn4^FlpO/FlpO^ animals also showed similar tau of the post-illumination pupil response (PIPR) compared to heterozygotes and controls (Figure 7F). However, the pupils of Opn4^FlpO/FlpO^ animals were significantly more dilated after 20 seconds in darkness following light offset (Figure 7E,H,J). Collectively, Opn4^FlpO/FlpO^ mice show similar deficits in the consensual PLR as other Opn4^−/−^, validating its functional use as an Opn4^−/−^ model for behavior.

**Figure 7.**
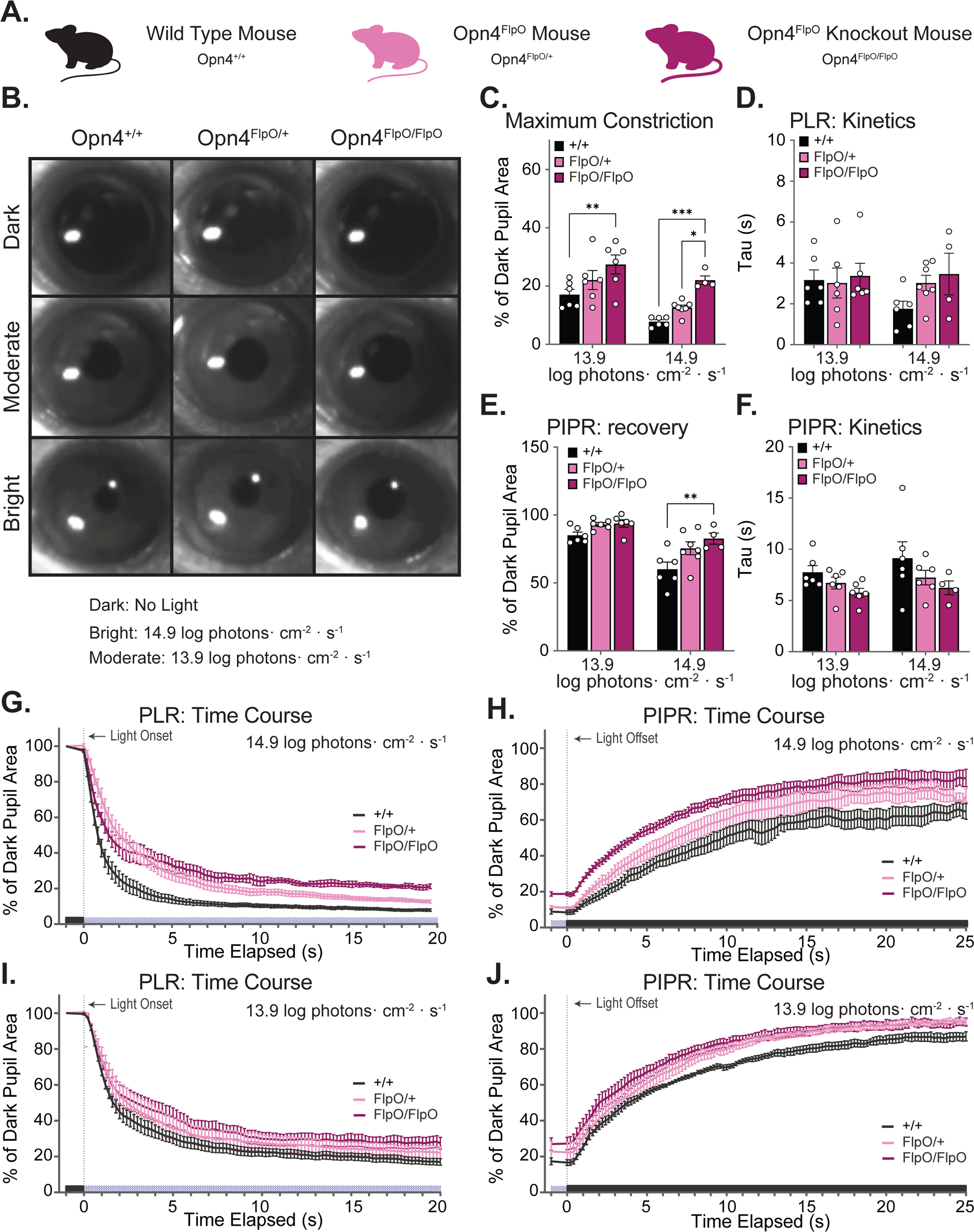
Opn4^FlpO/FlpO^ mice have deficits in the consensual pupillary light response reported in other Opn4^−/−^ mouse models. A. Schematic of three mouse lines used for behavior: Opn4^+/+^, Opn4^FlpO/+,^ and Opn4^FlpO/FlpO^. B. Representative images from video recordings of the consensual pupillary light response (PLR) in Opn4^+/+^, Opn4^FlpO/+,^ and Opn4^FlpO/FlpO^ mice in darkness, and at maximum constriction in moderate (13.9 log photons· cm^−2^ · s^−1^) and bright (14.9 log photons· cm^−2^ · s^−1^) 480nm light. C. Maximum constriction of the pupil after 20 seconds differed (2-way ANOVA, F(2,29)=14.38, p<0.0001). Opn4^FlpO/FlpO^ pupils constricted to 22% of dark pupil size in bright light, larger than Opn4^FlpO/+^ (p=0.0215, mean=12.56%) and controls (p=0.0007, mean=7.775%). Opn4^FlpO/FlpO^ constricted to 27.38% of dark pupil size in moderate light, larger than controls (p=0.0057, mean=17.02%), but similar to Opn4^FlpO/+^ (p=0.2083, mean=22.06%). D. Time constant (Tau) values for exponential decay functions fit to individual mouse PLR timecourse datasets (20 seconds of 0.2s interval relative pupil area values in light) were similar across light levels (linear mixed-effects, F(2,13)=1.088, p=0.3657). E. Post-illumination pupil recovery (PIPR) after 20 seconds of darkness, following PLR stimulus, as a percentage of dark pupil size, differed (F(1,29)=32.2, p<0.0001) with a greater pupil size in Opn4^FlpO/FlpO^ mice compared to controls (p=0.0081). F. Time constant (Tau) values for exponential decay functions fit to individual mouse PIPR time course datasets were similar across light levels (linear mixed-effects, (F(2,28)=0.1590, p=0.8538). G. Bright light PLR: Average pupil area during 0.2 second intervals of 14.9 log photons· cm^−2^ · s^−1^light stimulus. PLR of both Opn4^FlpO/+^ and Opn4^FlpO/FlpO^ was attenuated over time in response to bright light (2-way ANOVA, time x genotype interaction, F(198,1386)=5.115, p<0.000) H. Bright light PIPR: Average pupil area during 0.2 second intervals of darkness, after 14.9 log photons· cm^−2^ · s^−1^light stimulus. I. Moderate light PLR: Average pupil area during 0.2 second intervals of 13.9 log photons· cm^−2^ · s^−1^ light stimulus. PLR over time was similar between groups (2-way ANOVA, time x genotype, (F(202,1515)=0.6115, p=0.6110). J. Moderate light PIPR: Average pupil area during 0.2 second intervals of darkness, after 13.9 log photons· cm^−2^ · s^−1^ light stimulus. C-F, data points represent one mouse. G-J, mean +/− SEM

### Brain Projections of FlpO-positive RGCs

ipRGCs project widely to a defined subset of image- and non-image-forming visual brain regions. To determine whether the Opn4^FlpO^ allele can be used to trace downstream targets, we imaged projections from the retina in Ai65-F mice, or after intravitreal injection of a FlpO-dependent AAV reporter virus (AAV2/Ef1a-fDIO-mCherry) in adult retina.

Intravitreal AAV injection resulted in mCherry reporter expression both the cell bodies and axons of ipRGCs (Figure 8). mCherry reporter+ axons were visible in ipRGC-recipient areas including dorsal and ventral lateral geniculate nucleus (LGN), intergeniculate leaflet (IGL), olivary pretectal nucleus (OPN), the suprachiasmatic nucleus (SCN), lateral habenula (LHb), and superior colliculus (SC) (Figure 8) (Hattar et al., 2006; Baver et al., 2008; Güler et al., 2008, Ecker et al., 2010; Estevez et al., 2012; Li and Schmidt 2018). The SCN and OPN are primarily innervated by M1 ipRGCs while non-M1 ipRGCs innervate the SC (Hattar et al., 2006; Baver et al., 2008; Güler et al., 2008, Ecker et al., 2010; Estevez et al., 2012; Li and Schmidt 2018). As expected, the heaviest innervation was observed in non-image forming targets, consistent with our previous findings that most FlpO neurons are M1-M3 ipRGCs.

**Figure 8.**
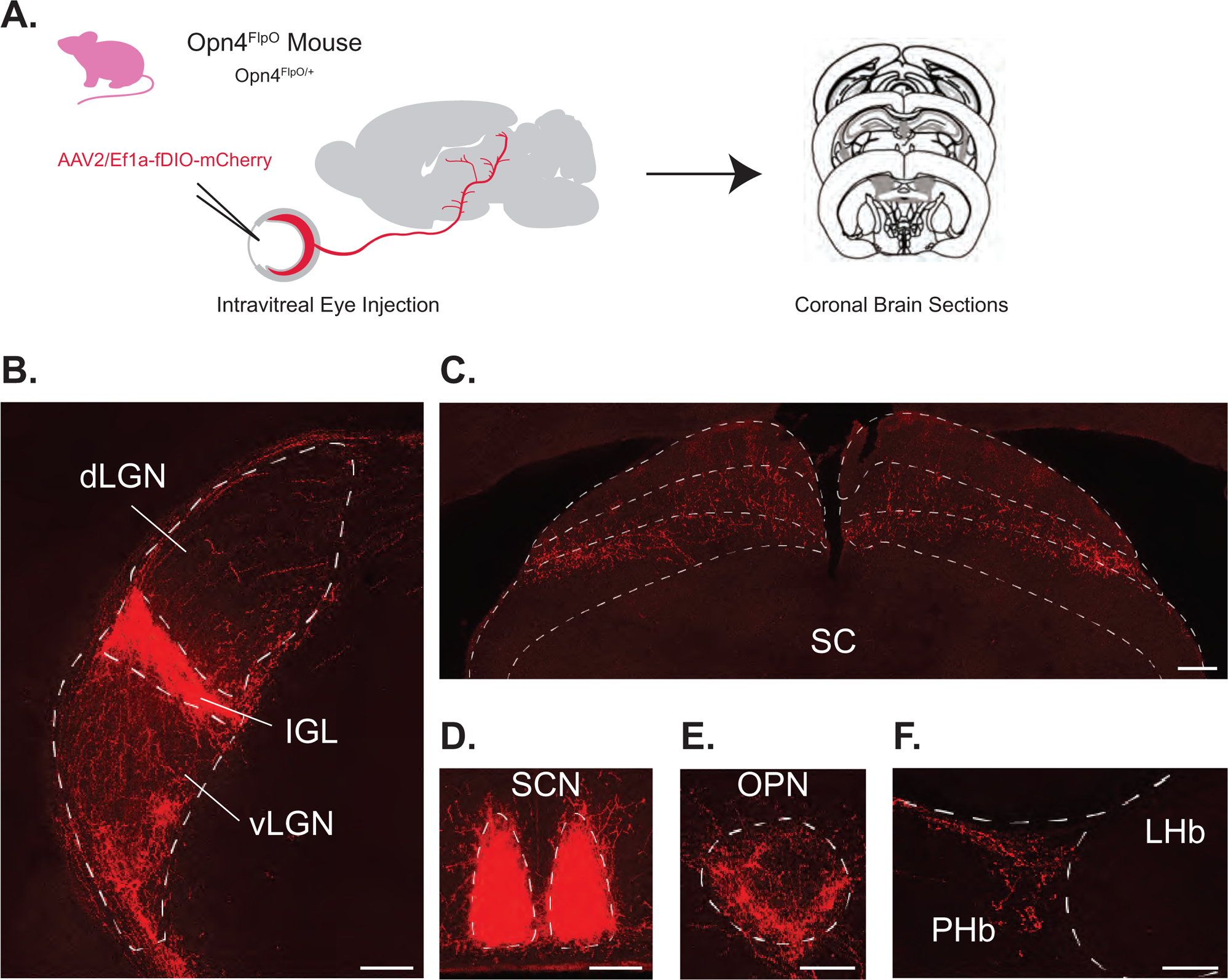
Retinofugal Projections in ipRGC-recipient regions using FlpO-dependent AAV reporter. A. AAV2-Ef1a-fDIO-mCherry was injected intravitreally into both eyes of adult Opn4^FlpO/+^ mice. B. – F. Representative coronal sections containing mCherry positive axons visualized in retinorecipient nuclei that receive input from ipRGCs: LGN: lateral geniculate nucleus (B), SC: superior colliculus (C), SCN: suprachiasmatic nucleus (D), OPN: olivary pretectal nucleus (E), and PHb: perihabenular region (F). Non-image-forming brain regions (IGL, vLGN, SCN, OPN) are more densely labeled than image-forming brain regions (dLGN and SC). Scale bars: 200 μm.

A similar pattern of labeling was also observed in the brains of Ai65-F (Opn4^FlpO/+^; Rosa26^Ai65-F/+^) mice, though the labeling was slightly more faint (Supporting Information Figure S8). In addition, we noted some extra-retinal labeling in brain neurons, as has been reported with Opn4^Cre^ animals (Ecker et al., 2010). Furthermore, we found no labeling in Rosa^Ai65-F/+^ animals that were not crossed to Opn4^FlpO^ animals (Supporting Information Figure S9). This indicates that all of the observed labeling is a result of Flp-dependent recombination in Ai65-F (Opn4^FlpO/+^; Rosa26^Ai65-F/+^) mice, even though the labeling was overall dimmer than in AAV labeled animals (Figure 8 and Supporting Information Figure S8). This suggests that, while both approaches may label or manipulate similar visual circuitry, AAV manipulations in adult animals may be more specific and may therefore be preferable depending on experimental design and availability of reagents.

## Discussion

In this paper we describe a new Opn4^FlpO^ line that drives expression of Flp-recombinase in ipRGCs. When Opn4^FlpO^ animals were crossed to multiple Flp-dependent reporter mouse lines, we consistently saw labeling in only RGCs as evidenced by the near complete overlap between fluorescent reporter labeling and RBPMS (Figure 2; Supporting Information Figure S4; Supporting Information Figure S5; Supporting Information Figure S6). Moreover, we saw colocalization with ipRGC markers and a lack of overlap with Brn3a, a transcription factor broadly expressed in non-ipRGCs (Figure 5; Supporting Information Figure S4; Supporting Information Figure S5; Supporting Information Figure S6). The total number of labeled RGCs per retina varied across each retina, with RCE::FrePe having the least and RCE:FRT having the most cells, with over 1000 cells in immersion fixed retinas and over 2000 cells labeled in perfusion-fixed animals (Figure 1; Supporting Information Figure S3). The vast majority of cells in all three lines co-labeled with Opn4, which labels the M1-M3 subtypes, indicating that this mouse line most effectively drives recombination in these ipRGC subtypes and may be a useful tool for manipulating these populations (Figure 3; Supporting Information Figure S4; Supporting Information Figure S5; Supporting Information Figure S6). This conclusion is supported by strong labeling of ipRGC axons in FlpO animals in M1-recipient targets such as the SCN (80% M1 and 20% M2) and IGL, along with visible, but sparser, labeling of non-M1 ipRGC targets such as the dLGN and SC, that would be innervated almost exclusively by non-M1 ipRGC subtypes (Figure 8; Supporting Information Figure S8) (Berson et al., 2002; Hattar et al., 2002, 2006; Güler et al., 2008; Hatori et al., 2008; Ecker et al., 2010; Estevez et al., 2012; Schmidt et al., 2014; Zhao et al., 2014; Stabio et al., 2018; Quattrochi et al., 2019; Sonoda et al., 2018; Huang et al., 2019). It is important to note that there was some labeling in the brains of animals crossed to Flp-dependent, lineage-tracing mouse lines that was absent when using intravitreal AAV transduction of FlpO-dependent constructs to fluorescently label ipRGCs (Figure 8; Supporting Information Figure S8). This indicates that some nonspecific brain labeling is present in the Opn4^FlpO^ line, similar to what has been reported for one Opn4^Cre^ line (Ecker et al., 2010). Nonetheless, in the retina, our data indicate that FlpO-dependent reporter lines label ipRGCs with high specificity.

One obvious beneficial use of the Opn4^FlpO^ line will be the use of intersectional approaches with Cre lines that may label subsets of RGCs that include ipRGCs, or with other Cre-expressing lines or vectors. The confinement of FlpO-driven reporter expression to mainly M1-M3 ipRGCs suggests that this mouse line is best suited for manipulation of these ipRGC subtypes. Notably, a small proportion of cells in Opn4^FlpO^ animals express SMI-32, suggesting that a very small subset of M4 ipRGCs are also labeled (Figure 4; Supporting Information Figure S4; Supporting Information Figure S5; Supporting Information Figure S6). This is an important consideration when designing future experiments using this mouse line. However, this feature could also be beneficial in allowing for intersectional manipulation of only a small subset of ipRGC subtypes, but the fidelity of this approach would of course have to be further validated on an experiment-by-experiment basis. The variation we observed across reporter lines shows that recombination efficiency is dependent on the reagents used. It is therefore important to note that, as with any genetic manipulation, recombination efficiency should be assessed for each experimental paradigm.

Functionally, the Opn4^FlpO/FlpO^ mouse line shows the expected deficits in pupil constriction in bright light and enhanced relaxation following light offset that are typical of melanopsin null animals (Figure 7). Thus, the line can be used in behavioral studies assessing not only ipRGC cellular and circuit function, but also of the role of melanopsin signaling in various light-evoked behaviors. Indeed, this FlpO line opens up the exciting possibility of new multi-recombinase crosses, including intersectional manipulations that could lead to new insights into the role of single ipRGC subtypes in light-evoked behavior.

---

**Figure S1.**
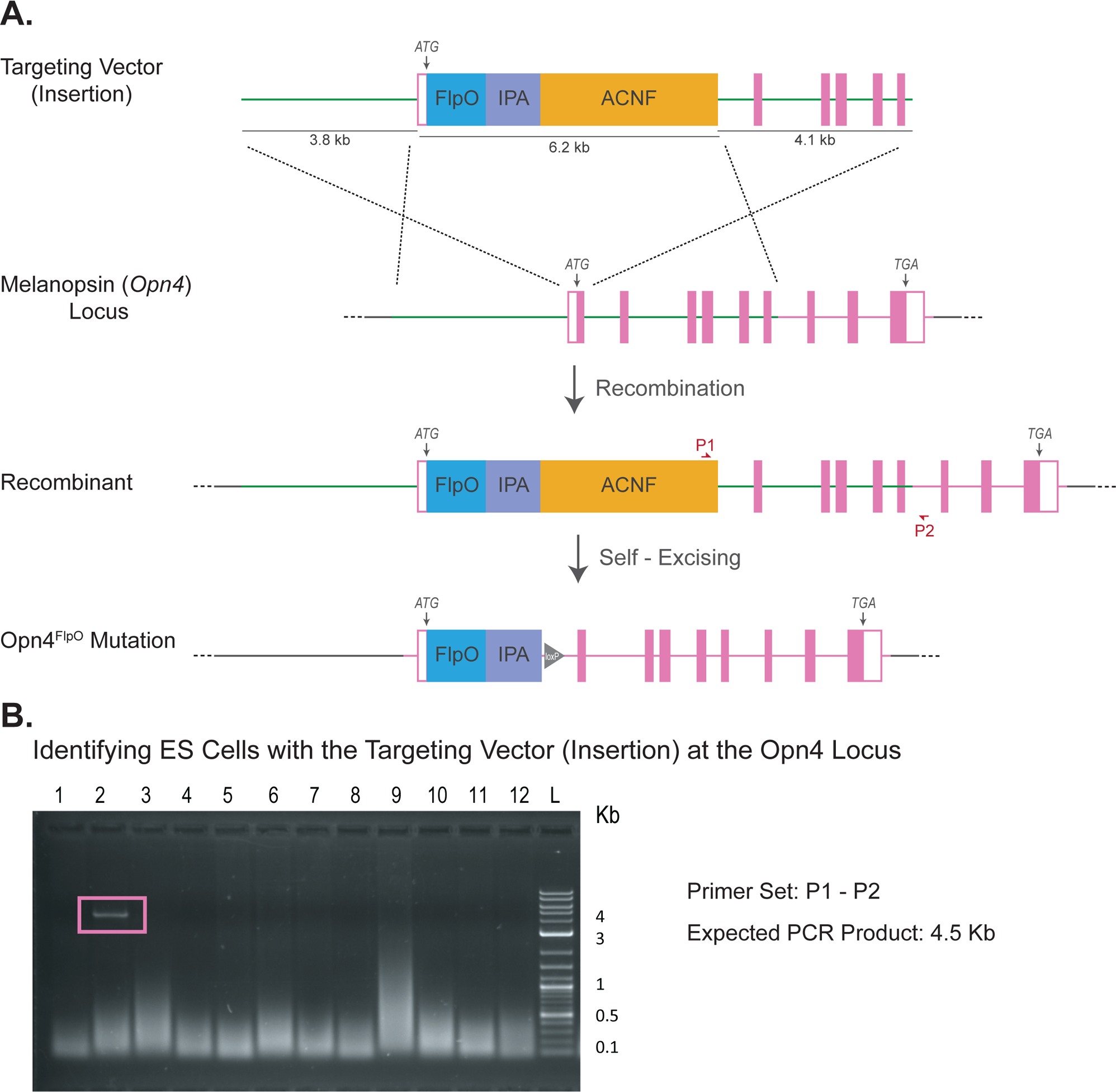
Generation of Opn4^FlpO^ mouse line. A. Top, diagram of the gene targeting strategy. The targeting vector contains a 6.2 kb insertion consisting of the FlpO coding sequence (blue box, FlpO) and an artificial intron and polyadenylation sequence (purple box, IPA), and an auto-excising neomycin selection cassette (orange box, ACNF), flanked by a 3.8 kb 5′ homology arm and a 4.1 kb 3′ homology arm. Homology arms are shown in green, and recombination sites indicated by black dotted lines. Middle, the inserted FlpO coding sequence is inserted flush with the endogenous Opn4 start codon (ATG). Primers used to screen for targeted insertion (P1 and P2) are indicated (red arrows). Bottom, a diagram of the Opn4^FlpO^ mutation that results after germline self-excision of the ACNF cassette, leaving behind a single loxP site. B. Agarose gel electrophoresis of the 4.5 kb PCR product generated from long-range PCR amplification. Primer Set P1 – P3 was designed to produce a PCR product outside the targeting vector to screen ES cell clones with the targeting vector insertion at the melanopsin locus in the correct orientation. Lanes 1-12 are the same individual ES cell clones shown in (B). The pink box is around the PCR amplicon in lane 2 from an ES cell clone contained the targeting vector in the correct place and orientation. Lane L is the 1 kb DNA size marker.

**Figure S2.**
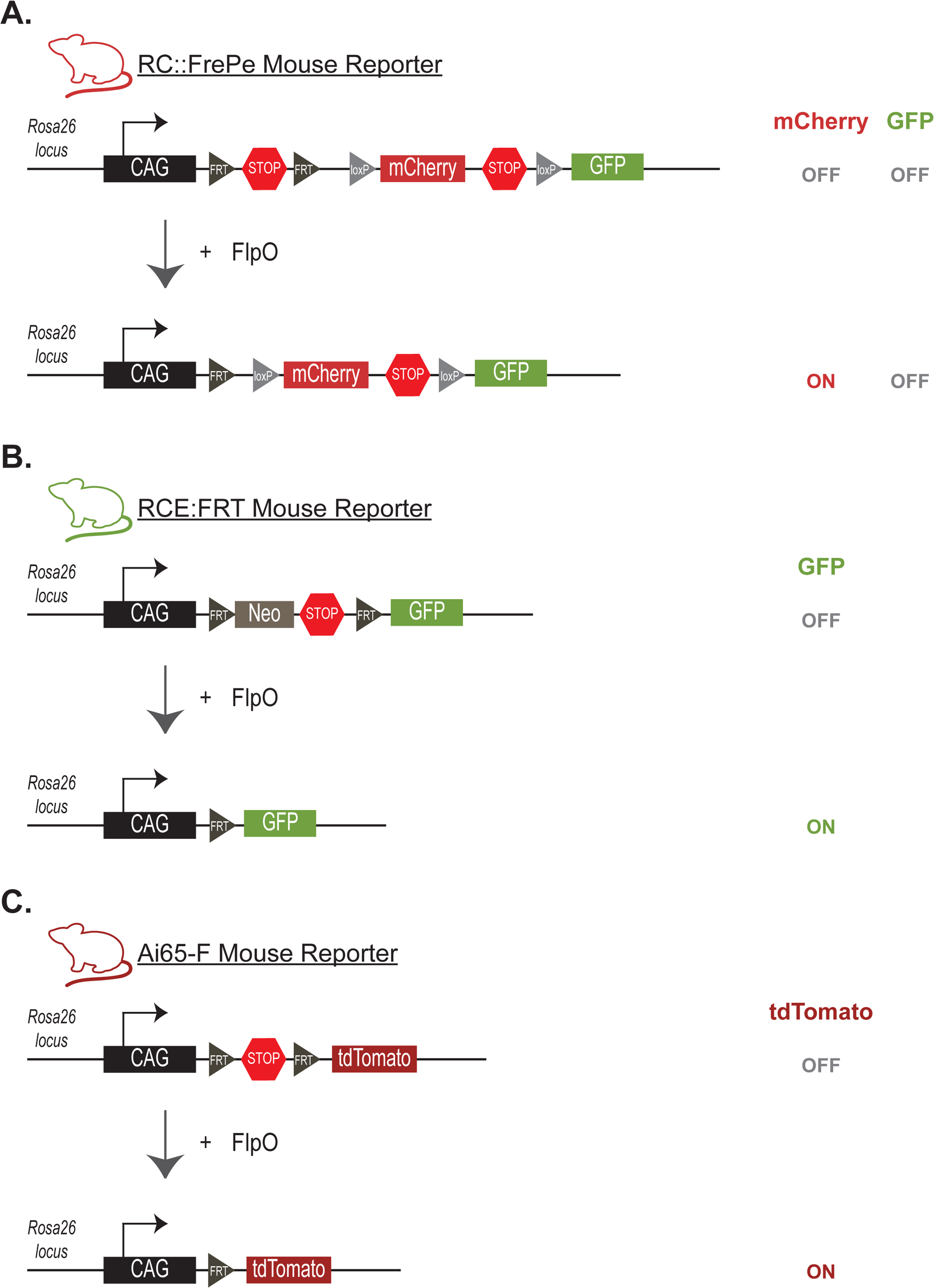
FlpO dependent reporters crossed to Opn4^FlpO^. A. The RC::FrePe mouse reporter utilizes a CAG (CMV (cytomegalovirus) early enhancer/chicken β-actin) promoter in the ROSA26 locus to drive gene reporter expression. RC::FrePe is a dual recombinase reporter with a STOP sequence flanked by FRT Flp recombinase recognition sites followed by an mCherry sequence and a STOP sequence flanked by loxP Cre recombinase recognition sites. FlpO recombinase drives expression of mCherry but expression of GFP is dependent on the presence of both Cre and FlpO recombinases (Bang et al., 2012). B. The RCE:FRT mouse reporter utilizes a CAG promoter in the ROSA26 locus with a neomycin and STOP sequence flanked by FRT sites. The single recombinase reporter drives expression of GFP in the presence of FlpO recombinase (Sousa et al., 2009). C. The Ai-65F mouse reporter utilizes a CAG promoter in the ROSA26 locus with a STOP sequence flanked by FRT sites. FlpO recombinase drives expression of tdTomato in the single recombinase reporter Ai-65F (Daigle et al., 2018).

**Figure S3.**
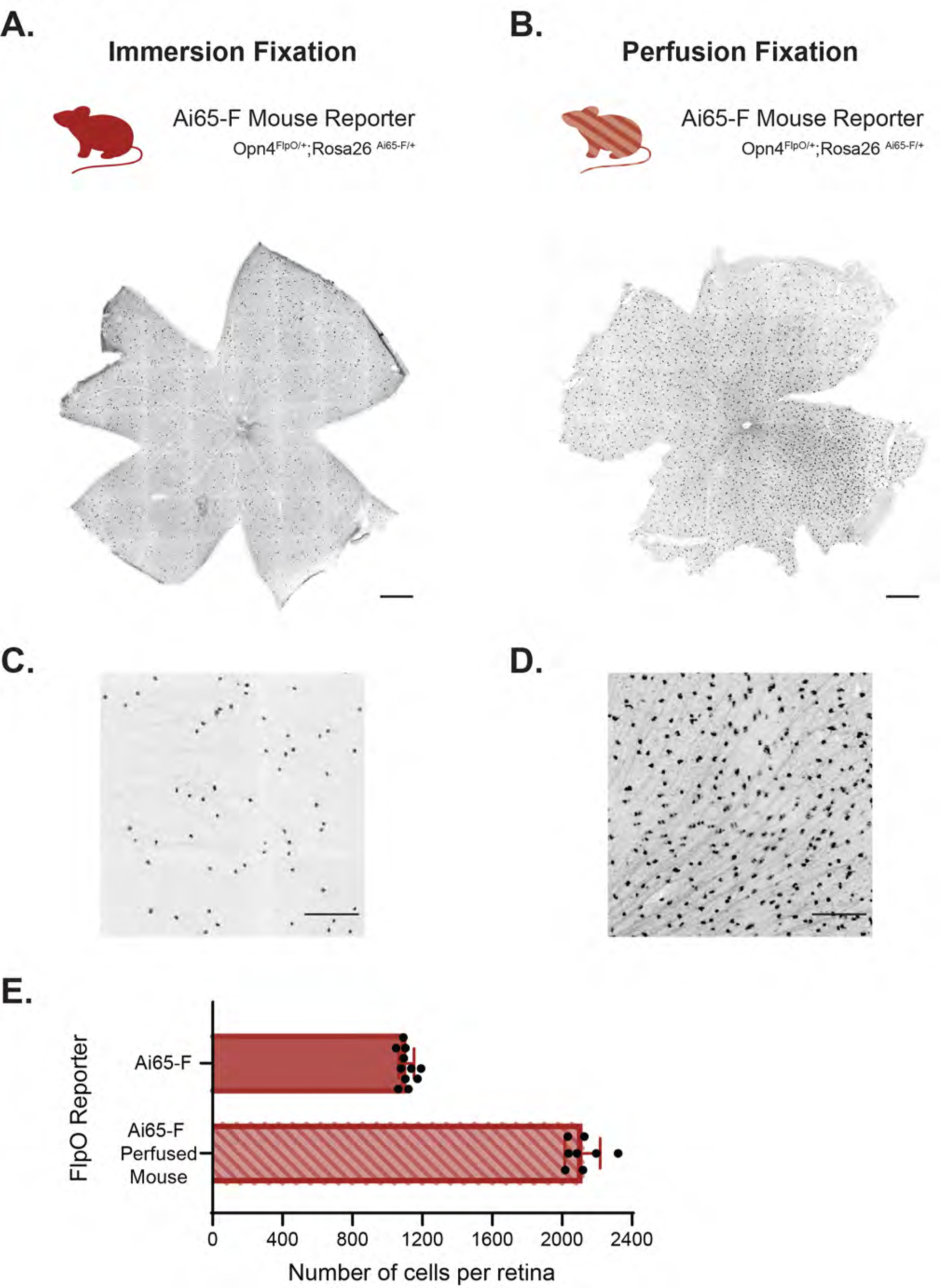
Perfusion fixed Ai65-F retinas contain twice as many cells as immersion fixed Ai65-F retina. A. Representative image of a whole-mount retina from an Ai65-F (Opn4^FlpO/+^;Rosa26^Ai65-F/+^) mouse processed via immersion fixation. This image is shown in Figure 1B, right side panel. Scale bar 500μm. B. Representative image of a whole-mount retina from an Ai65-F (Opn4^FlpO/+^;Rosa26^Ai65-F/+^) animal processed via transcardial perfusion fixation. The retina shows dense cell labeling compared to the retina that was immersion-fixed. Scale bar 500μm. C. High magnification image from the retina shown in (A). Image is shown in Figure 1C, right side panel. Scale bar 200μm. D. High magnification image from the retina shown in (B). The perfusion fixed retina labels more cells compared to the immersion-fixed retina in (C) Scale bar 200μm. E. Graph compares the total number of cells per retina from perfusion fixed retinas and immersion-fixed retinas isolated from Ai65-F (Opn4^FlpO/+^;Rosa26^Ai65-F/+^) animals. Individual dots on the graph represent the number of labeled cells in one retina. The number of labeled cells per retina was obtained by assessing cell labeling in the ganglion cell layer and the inner nuclear layer. Immersion-fixed had 1109 ± 44 labeled cells per retina (n=11 retinas from 7 mice) compared to 2117 ± 101 labeled cells per retina (n=8 retinas from 4 mice) from perfused fixed retinas isolated from Ai65-F animals. On average, perfused fix

**Figure S4.**
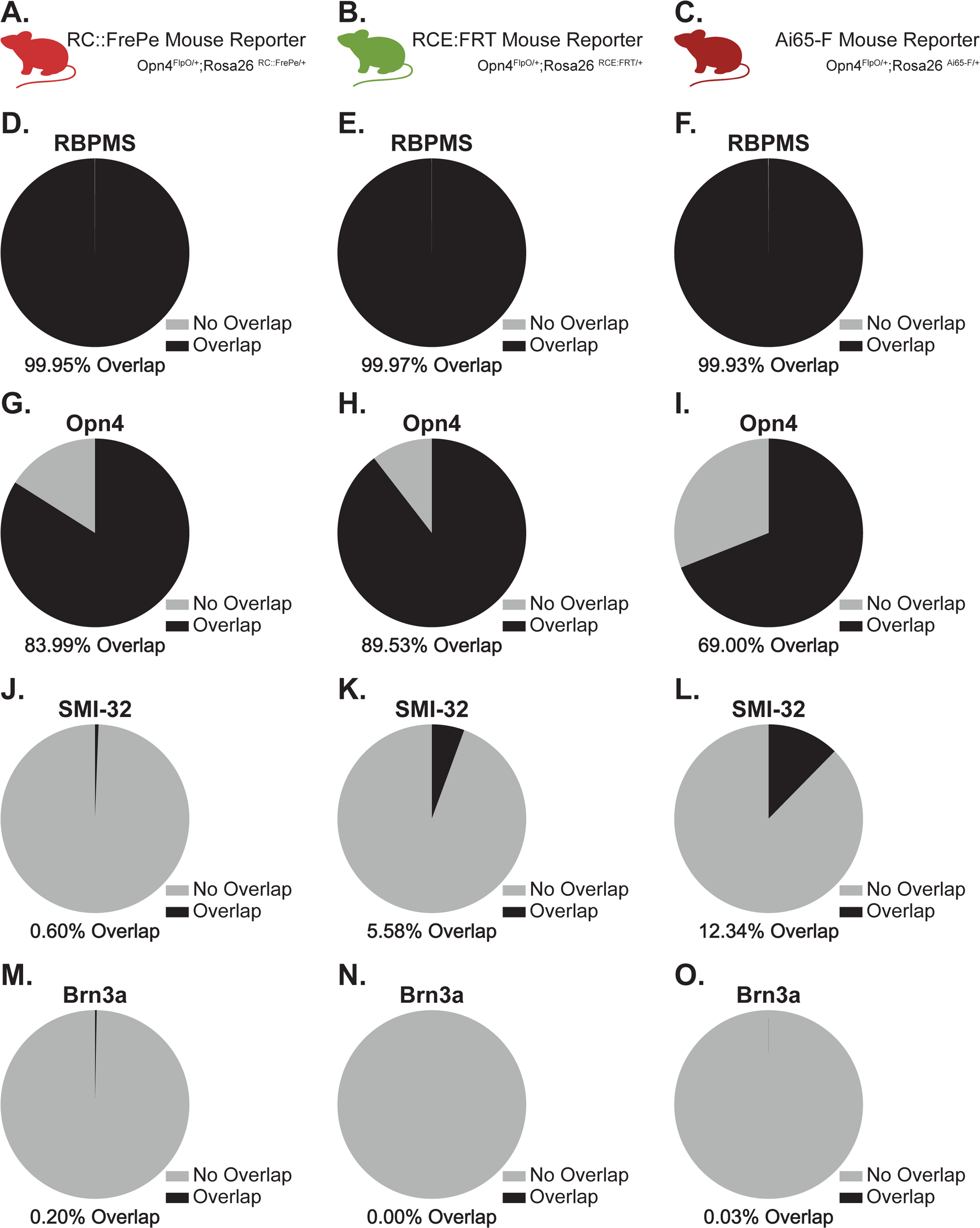
Similar proportion of co-labeling of fluorescent reporter with different markers across multiple FlpO dependent reporters. A. – C. Schematic of three FlpO dependent reporters: RC::FrePe, RCE:FRT, and Ai65-F used to characterized the Opn4^FlpO^ mouse line. D. – F. Proportion of fluorescent reporter positive cells labeled with RBPMS for all three lines. More than 99% of cells expressing the fluorescent reporter co-labeled with RBPMS in each reporter line. G. – I. Proportion of fluorescent reporter positive cells labeled with Opn4 for all three lines. The majority of cells expressing the fluorescent reporter are co-labeled with Opn4 and likely M1-M3 ipRGCs. J. – L. Proportion of fluorescent reporter positive cells labeled with SMI-32 for all three lines. Proportion of SMI-32 differed slightly across reporter lines. M. – N. Proportion of fluorescent reporter positive cells labeled with Brn3a, very few cells showed colocalization of fluorescent reporter expression and Brn3a labeling.

**Figure S5.**
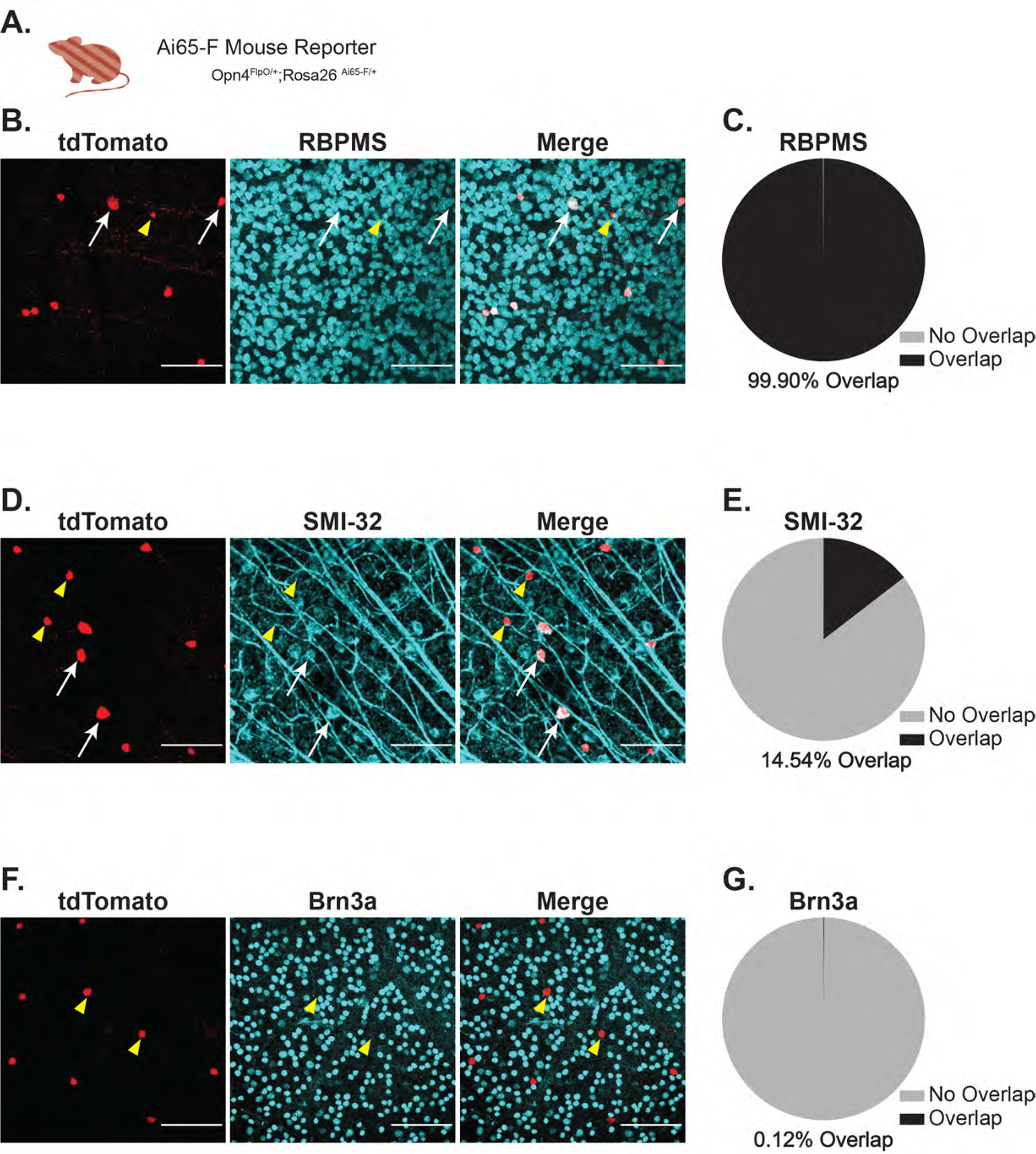
Proportion of cell markers labeled in perfused fixed retinas. A. Retinas from Ai65-F (Opn4^FlpO/+^;Rosa26^Ai65-F/+^) animals were fixed via transcardial perfusion with 4% paraformaldehyde. B. Example images of the GCL in a whole-mount retina immunolabeled for tdTomato (red, left panel) and RBPMS (cyan, middle panel). Right panel shows merged image. C. Total proportion of tdTomato cells co-labeled (black) or not (gray) with RBPMS (n=4 retinas from 2 mice). D. Representative Images of the GCL in a whole-mount retina immunostained for tdTomato (red, left panel) and SMI-32 (cyan, middle panel). Right panel shows merged image. E. Total proportion of tdTomato positive cells co-labeled (black) or not (gray) with SMI-32 (n=4 retinas from 2 mice). F. Representative images of the GCL in a whole-mount retina immunolabeled for tdTomato (red, left panel) and Brn3a (cyan, middle panel). Right panel shows merged image. G. Total proportion of tdTomato positive cells co-labeled (black) or not (gray) with Brn3a (n=2 retinas from 1 mice). White arrows denote fluorescent reporter positive cells co-expressing the cell marker. Yellow triangles illustrate reporter cells lacking overlap with the cell marker. Scale bar 100μm.

**Figure S6.**
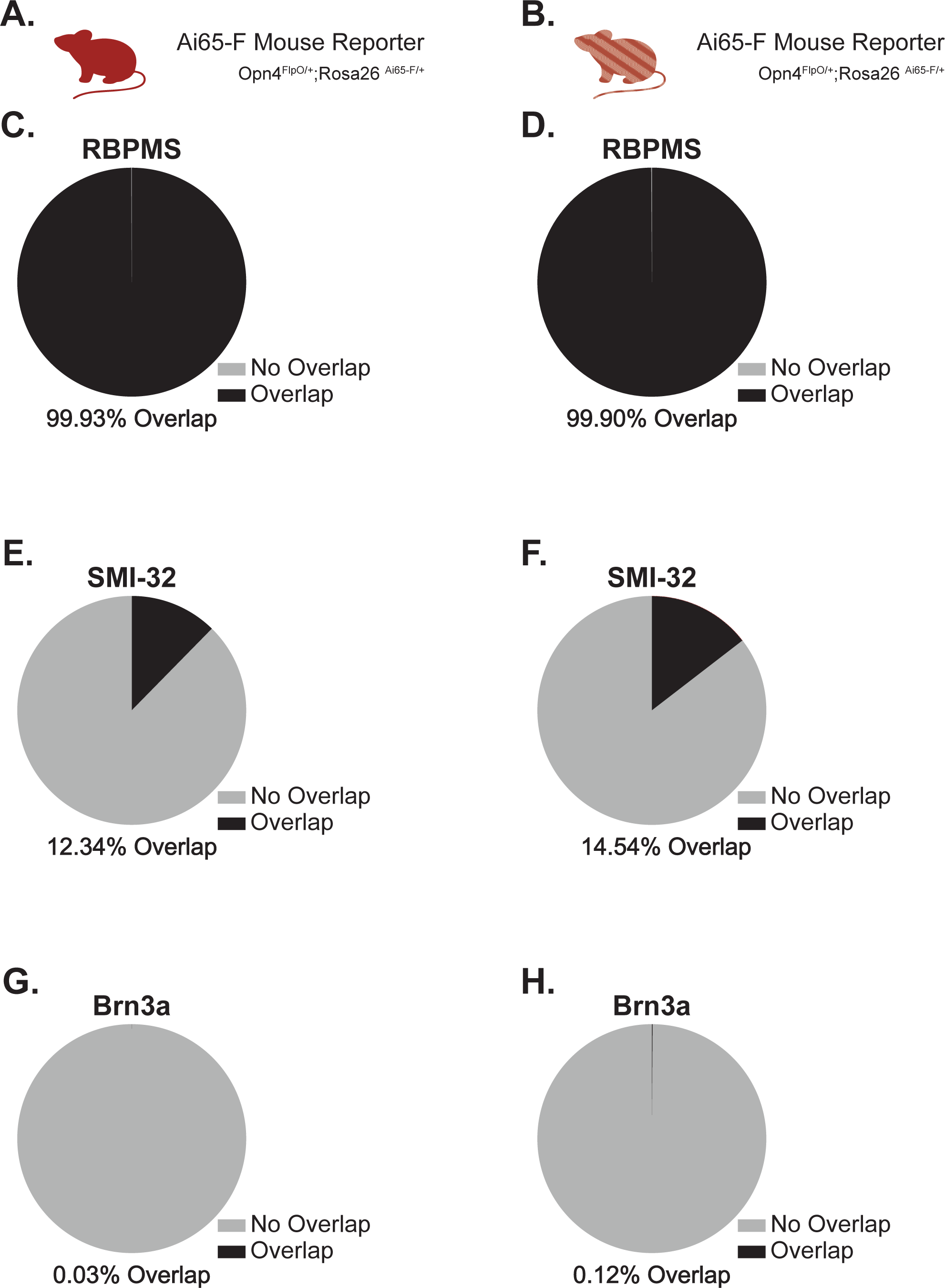
Immersion and perfused fixed Ai65-F retinas show similar proportions of labeled cell markers. A. Retinas from Ai65-F (Opn4^FlpO/+^;Rosa26^Ai65-F/+^) animals were fixed via immersion fixation with 4% paraformaldehyde. B. Retinas from Ai65-F (Opn4^FlpO/+^;Rosa26^Ai65-F/+^) animals were fixed via transcardial perfusion with 4% paraformaldehyde. C. Total proportion of tdTomato positive cells co-labeled (black) or not (gray) with RBPMS in immersion fixed retinas (n=5 retinas from 3 mice). D. Total proportion of tdTomato positive cells co-labeled (black) or not (gray) RBPMS in perfusion fixed retinas (n=4 retinas from 2 mice). E. Total proportion of tdTomato positive cells co-labeled (black) or not (gray) with SMI-32 in immersion fixed retinas (n=4 retinas from 2 mice). F. Total proportion of tdTomato positive cells co-labeled (black) or not (gray) with SMI-32 in perfusion fixed retinas (n=4 retinas from 2 mice). G. Total proportion of tdTomato positive cells co-labeled (black) or not (gray) with Brn3a in immersion fixed retinas (n=3 retinas from 2 mice). H. Total proportion of tdTomato positive cells co-labeld (black) or not (gray) with Brn3a in perfusion fixed retinas (n=2 retinas from 1 mice). From Supporting Information Figure S5G. F. ed retinas label almost double the number of cells than immersion-fixed retinas.

**Figure S7.**
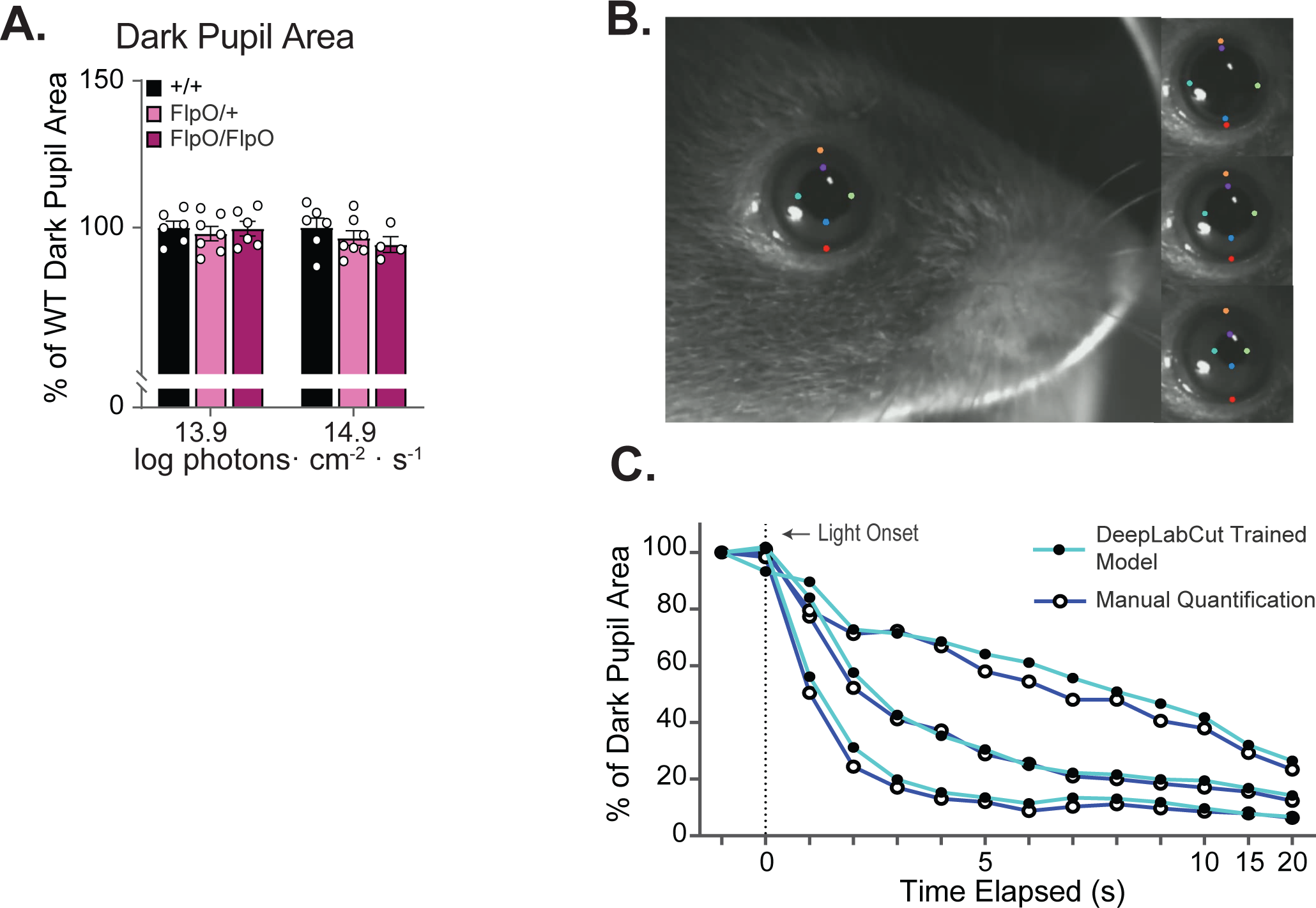
DeepLabCut pupil tracking and validation. A. Pupil areas were equal in darkness (2-way ANOVA, F(2,14)=0.6063, p=0.5591) Dark pupil area calculated as a percentage of eye area, and averaged across the 120 frames (2 seconds) of video immediately prior to light stimulus, then normalized to control. B. Representative images from a video analyzed and labeled using a machine learning model trained for 350k iterations in DeeplabCut. Innermost tracked points (violet and blue, green and mint) were used to calculate the area of the pupil. Outermost tracked points (orange and red) were used to normalize the pupil area to the size of the eye and correct for small changes in the distance between the eye and the camera. C. Three randomly chosen videos were manually quantified by measuring the pupil and eye diameter using the line tool in Fiji, from which the relative pupil area was calculated in 20 frames per video, one per second (navy). There was no significant difference between these, and pupil areas calculated from DeepLabCut generated coordinates (cyan). (2-way ANOVA, time x quantification F(13, 52)=0.06946, p>0.9999, quantification F(1,4)=0.03253, p=0.8656)

**Figure S8.**
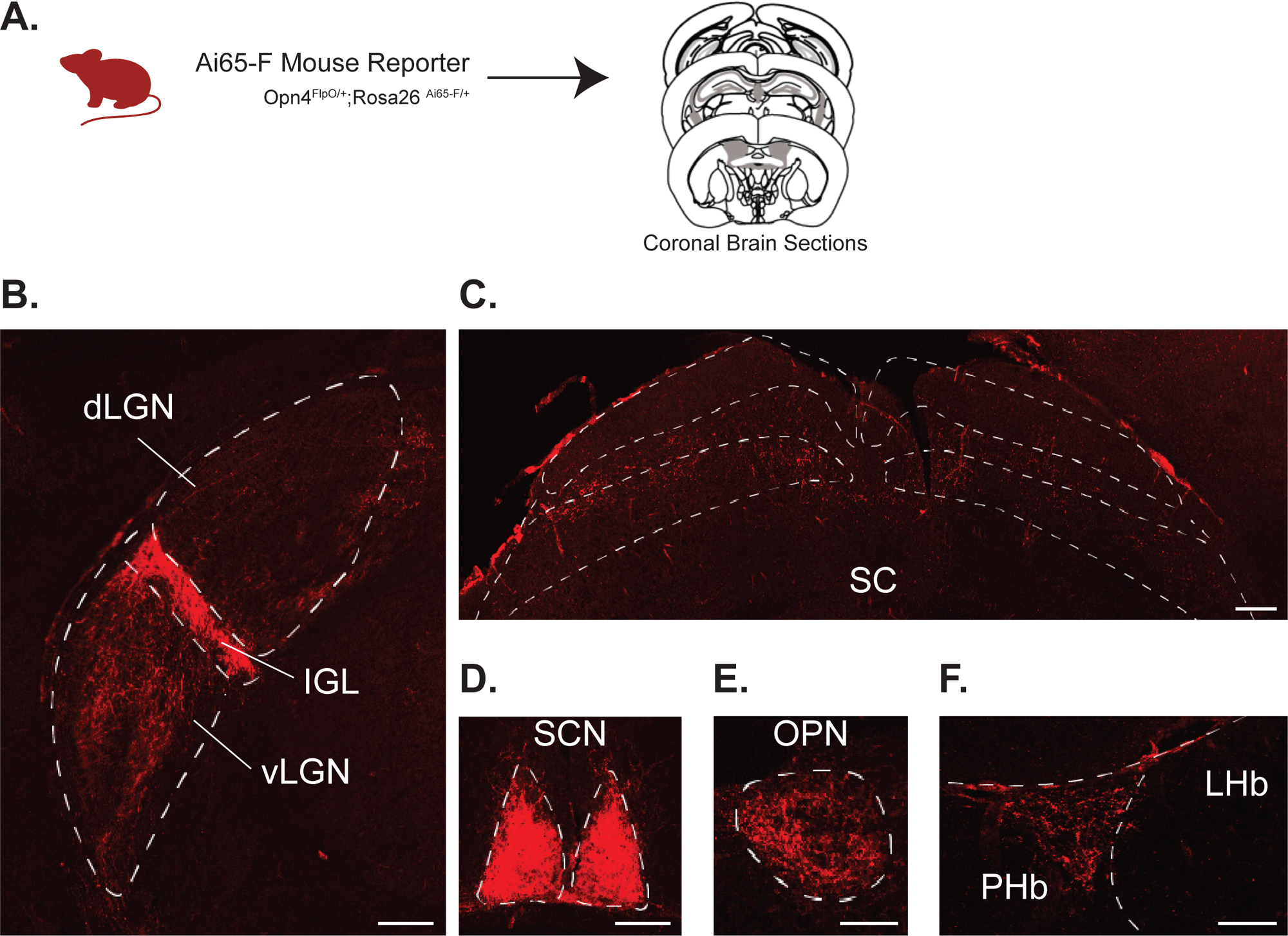
Retinofugal Projections in ipRGC-recipient regions using Ai65-F reporter line. A. Coronal sections were taken from Ai65-F (Opn4^FlpO/+^; Rosa^Ai65-F/+^)animals. B. – F. Representative coronal sections containing tdTomato positive axons visualized in retinorecipient nuclei that receive input from ipRGCs: LGN: lateral geniculate nucleus (B), SC: superior colliculus (C), SCN: suprachiasmatic nucleus (D), OPN: olivary pretectal nucleus (E), and PHb: perihabenular region (F). Similar to AAV reporter labeling, non-image-forming brain regions (IGL, vLGN, SCN, OPN) are more densely labeled than than image-forming brain regions (dLGN and SC). Sparse ectopic labeling was observed in some brain neurons. Scale bars: 200 μm.

**Figure S9.**
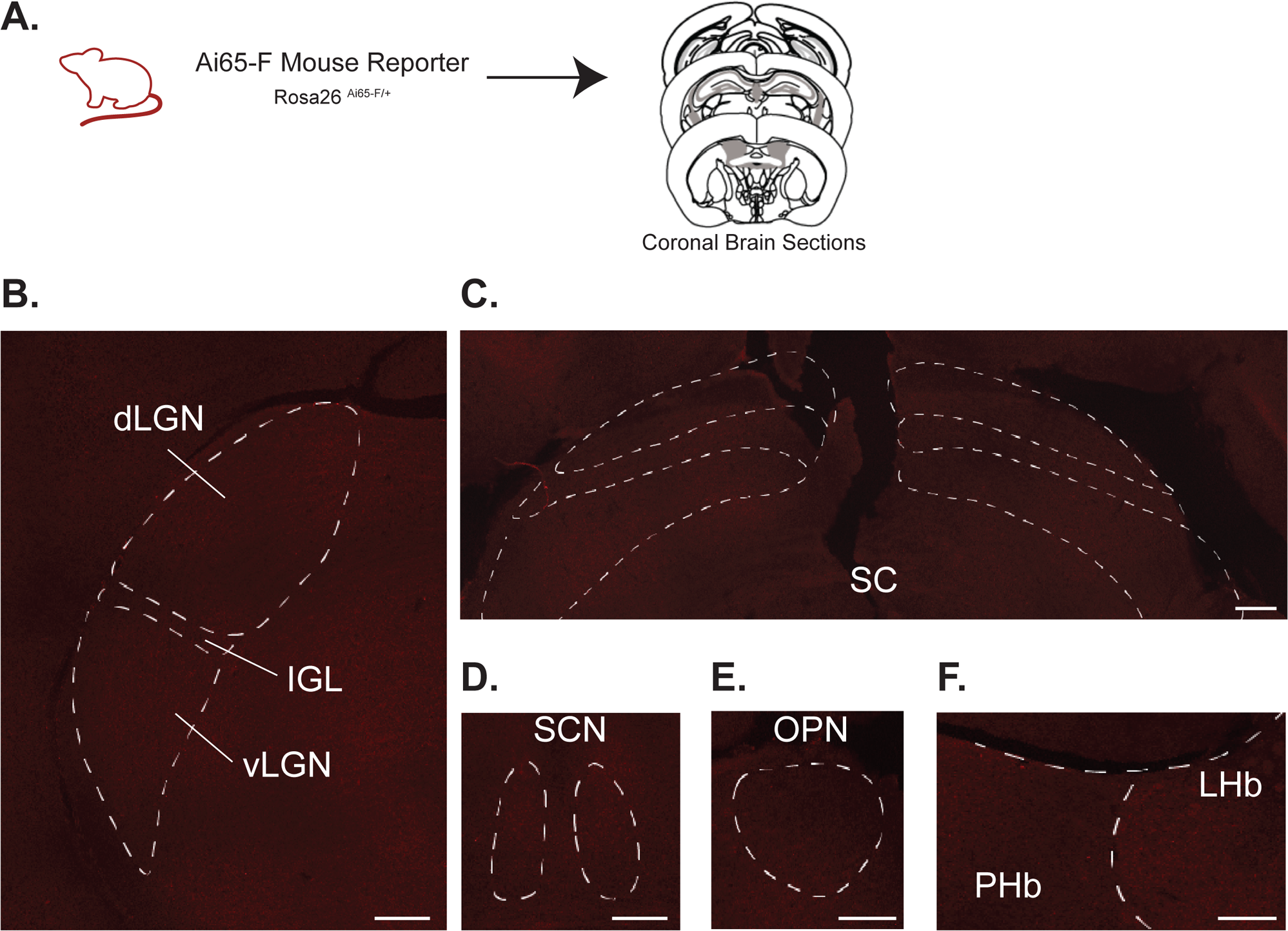
No labeling observed in Opn4^+/+^; Rosa^Ai65-F/+^ reporter mice lacking FlpO expression. A. Coronal sections were taken from Opn4^+/+^; Rosa^Ai65-F/+^ reporter mice lacking FlpO. B. – F. No tdTomato axons were observed in ipRGC-recipient regions shown in Figure 8 and Supporting Information Figure S8: LGN: lateral geniculate nucleus (B), SC: superior colliculus (C), SCN: suprachiasmatic nucleus (D), OPN: olivary pretectal nucleus (E), and PHb: perihabenular region (F). Scale bars: 200 μm.

## Acknowledgements

We thank Amanda Menzie and Hardik Patel for technical support in generating the mouse strain, and Lynn Doglio and the Transgenesis and Targeted Mutagenesis Laboratory at Northwestern University for ES gene targeting and chimera production.

